# Interspecific relationships in a suboptimal habitat, an unexpected ally for the survival of the threatened Tenerife speckled lizard (*Gallotia intermedia*)

**DOI:** 10.1101/2023.12.11.571074

**Authors:** Gonzalo Albaladejo-Robles, Alejandro Escánez, Alicia V. Perera-Castro

## Abstract

Anthropogenic-driven environmental changes are pushing species to the limits of their habitats. More often species are restricted to relic or suboptimal habitats that present the minimum requirements to sustain species populations. In this scenario of accelerated environmental change and biodiversity loss, is fundamental to understand why species can survive in such suboptimal conditions. We conduct an isotopic trophic analysis along with a behavioural experiment to show how novel ecological interactions allow an endangered species to maintain stable populations in suboptimal habitats. We show how the Tenerife speckled lizard (*Gallotia intermedia*), a critically endangered endemic reptile from Tenerife Island (Canary Islands), can maintain stable populations in relic habitats thanks to its interactions with the yellow-legged gull (*Larus michahellis*) colony. A stable isotope analysis revealed that *G. intermedia* relies on marine subsidies for its diet and that the foraging area of this reptile is likely to be restricted to the limits of *L. michahellis* breeding colony. Furthermore, an antipredator behaviour analysis showed that *L. michahellis* displayed a strong anti-predator or mobbing response against cats, one of the main threats for *G. intermedia*, thus potentially providing some protection to the reptiles inhabiting the seabird colony. Our results show how unusual and poorly studied biotic interactions can provide valuable resources and conditions for the conservation of a critically endangered species inhabiting a suboptimal or relict habitat.

## Introduction

Biodiversity is facing a human-driven crisis. The last Living Planet Report showed an overall decrease of 68% in the abundance of monitored populations of vertebrates at a global scale (Dirzo et al., 2014; WWF, 2020). Globally, the main anthropogenic threats to biodiversity are habitat change, followed by overexploitation of species, climate change, pollution, and biological invasions (Brondizio et al., 2019; Dirzo et al., 2014). In many cases, these and other stressors are pushing species to the brink of extinction, with many species and populations forced to inhabit the edge of their distributions or refugial habitats, persisting over time in the face of biophysical disturbances (Keppel et al., 2012; Sedell et al., 1990). In many cases, these refuge habitats are considered suboptimal or marginal habitats because they do not provide the best habitat but the required environmental conditions (abiotic and abiotic context) for the species to reach a minimum fitness to fulfil their life histories, and in the case of refugee species this depictures their realized or contemporary niche (Brown & Carnaval, 2019; Kerley et al., 2020; Scheele et al., 2017). However, these marginal habitats may not be the most favourable and represent the least affected or most resilient habitat to the drivers of decline (Kerley et al., 2020). In contrast, populations located in these marginal environments may be less tolerant to increasing threat impacts (Scheele et al., 2017), which can turn these refuges into potential ecological traps (Robertson & Hutto, 2006). Even though the importance of these refugees is well recognized for biodiversity conservation (Braidwood et al., 2018; Keppel et al., 2015; Lea et al., 2016; Pavey et al., 2017; Selwood & Zimmer, 2020), it is still unclear, in most cases, which are the mechanisms and processes that allow species and populations to persist in these habitats, especially those involving biotic conditions (Gaston & Gaston, 2003).

Refugee species are those considered to have lost access to their past optimal habitats and are, therefore, currently confined to suboptimal habitats. Usually, these species follow three ‘rules of thumb’ 1) have suffered a severe historical range contraction, 2) maintain slow population growth rates and/or low densities despite conservation efforts, and 3) appear to make use of anomalous resources (e.g., dietary shifts) in the currently occupied habitat compared to historical records and that of close relatives (Kerley et al., 2012). Many threatened faunal species are recognized as refugee species (e.g., Mediterranean monk seal, European bison, kakapo or giant panda) in their current distributional areas (Kerley et al., 2020).

Biotic conditions or biotic environments, such as interspecific interactions (i.e., mutualism, commensalism, predation, parasitism, competition, etc.), in addition to availability and dynamic of resources, can generate positive interactions and boost species distribution and abundance in otherwise unsuitable areas (Godsoe et al., 2017; Soberón, 2007). Despite this, the relevance of these biotic interactions in shaping the large-scale distribution of a species is still a matter of debate (Godsoe et al., 2017; Peterson et al., 2011; Wiens, 2011). However, their effects at local scales on species ranges seem to be clear when barriers to its dispersal and abiotic gradients are weak (Davis et al., 1998; Guisan & Thuiller, 2005; Louthan et al., 2015; Wisz et al., 2013).

As a case study on the importance of interspecific interactions in refugee species living in a suboptimal habitat, we focus our work on the endemic Canarian giant lizards which are currently represented by four species of the same genus: *Gallotia simonyi* from El Hierro, *Gallotia bravoana*, from La Gomera and *Gallotia intermedia* from Tenerife. The Giant lizards of the Canaries are a good example of species widely distributed in the past, that have seen their natural distribution drastically reduced (Gonzalez et al., 2014; Palacios-García et al., 2021). Human activities, such as habitat destruction and especially the introduction of invasive alien species (IAS) such as cats and rats, have driven all three species to the brink of (Afonso & Mateo, 2009; Bonnaud et al., 2011; García Márquez et al., 1997; Medina & Nogales, 2009). Nowadays these “giants” are restricted to small populations of refugees in the most inaccessible and almost barren coastal cliffs of their respective islands (Nogales et al., 2001; J. C. Rando et al., 2000; J.-C. Rando et al., 2004). Due to all those factors, all the giant lizards from the Canary Islands are classified as Critically Endangered (CR) by the IUCN Red-list of Threatened Species and protected under Spanish and international legislation. From these species, only the Tenerife speckled lizard, *G. intermedia*, presents two natural populations in its native range, Teno massif and Guaza cliffs. This makes this species a candidate for refugee species and a perfect species to study which are the conditions that allow this group of “giant” lizards to maintain their population in suboptimal habitats.

Here we focus on the *G. intermedia* population located in Guaza mountain cliffs, which was discovered in 2003 (J.-C. Rando et al., 2004). Soon after its discovery, this species was classified as Critically endangered by the IUCN (Mateo et al., 2019), and Endangered by the Spanish legislation (BOC, 2010; BOE, 2011). This population is restricted to inaccessible platforms and some ravines along the coastal cliffs of Guaza Mountain, located in the south-west of Tenerife Island, where it shares space with one of the largest resident colonies of yellow-legged gull (*Larus michahellis*) of the island (Figure 1) (Albaladejo et al., 2015; J.-C. Rando et al., 2004, p. 20). Although this population was described more than two decades after the elaboration of this manuscript, little is known about the biotic interactions between *G. intermedia* and its immediate surrounding environment. As stated before, Guaza cliffs are poor in vegetation cover. This alone is a challenge for *G. intermedia*. All *Gallotia* genus lizards are mostly herbivorous, feeding on fruits, leaves, and flowers of a wide variety of plants but also include arthropods and other small animals, including carrion (Fariña & Martín, 2013; Machado, 1985; Pérez-Mellado et al., 1999; Rica, 1981). However, the presence of a stable colony *L. michahellis* creates a potential source of food for *G. intermedia*. Previous studies report observations of *G. simonyi* and *G. intermedia* feeding on the remains left by *L. michahellis* (Mateo, 2006; Siverio & Felipe, 2009). Furthermore, some studies show how seabird-derived resources are an important energy income for a variety of reptilian populations in areas with low primary productivity (Barret W.B. Anderson, D. A. Wait, L.L. Grismer, G.A. Polis & M.D. Rose et al., 2005; Kenny et al., 2017; Spiller et al., 2010). Therefore, we hypothesized that, in the absence of a substantial vegetation cover, *G. intermedia* is using seabird-derived resources left in the area by the permanent colony of *L. michahellis* as a complementary source of energy. Additionally, we believe that the interaction with *L. michahellis* goes beyond marine subsidies, acting also as indirect “bodyguards” or “protector species” for the reptiles (Gameiro et al., 2022). The abrupt and hard topography of Guaza coastal cliffs serves as a natural barrier for feral cats. Feral cats are the main responsible for the decline of several giant endemic lizards (genus Gallotia) in the Canaries, such as Gallotia simonyi (García Márquez et al., 1997), Gallotia intermedia (Albaladejo et al., 2015; Hernández et al., 2000; J.-C. Rando et al., 2004) and Gallotia gomerana (Nogales et al., 2001; J. C. Rando et al., 2000). The relatively low permeability of Guaza cliffs to this predator allows *G. intermedia* to inhabit the steepest profiles of these habitats with a lower risk of predation. However, as stated before, *G. intermedia* also uses the numerous platforms and some of the ravines present along the coastline. These areas are more permeable to cats, and so evidence of feral cat attacks and predation over *G. intermedia* have been recorded in those areas (Albaladejo et al., 2015; Flores Ravelo & Rando Reyes, 2021). These same ravines and platforms are used by *L. michahellis* colony for breeding and resting. The yellow-legged gull, like many other bird species, presents an aggressive anti-predator behaviour especially during the breeding season (Clode et al., 2000; Frixione & Salvadeo, 2021; Gameiro et al., 2022; Guidos et al., 2023). Previous studies showed how seabird breeding colonies, due to their aggressive response to predators, work as facilitators for the breeding of less aggressive and/or solitary species (Erwin, 1988; Gameiro et al., 2022; Quinn & Ueta, 2008; D. S. Richardson & Bolen, 1999). We hypothesize that *L. michahellis* anti-predator behaviour could also work in favour of *G. Intermedia*, providing populations inhabiting within the breeding grounds of this species some indirect protection against common predators. All these factors might explain, at least in part, the actual constrained distribution and conservation status of *G. intermedia* and to a larger extent the distribution of the rest of the giant reptiles of the Canary Islands.

**Figure 1.**
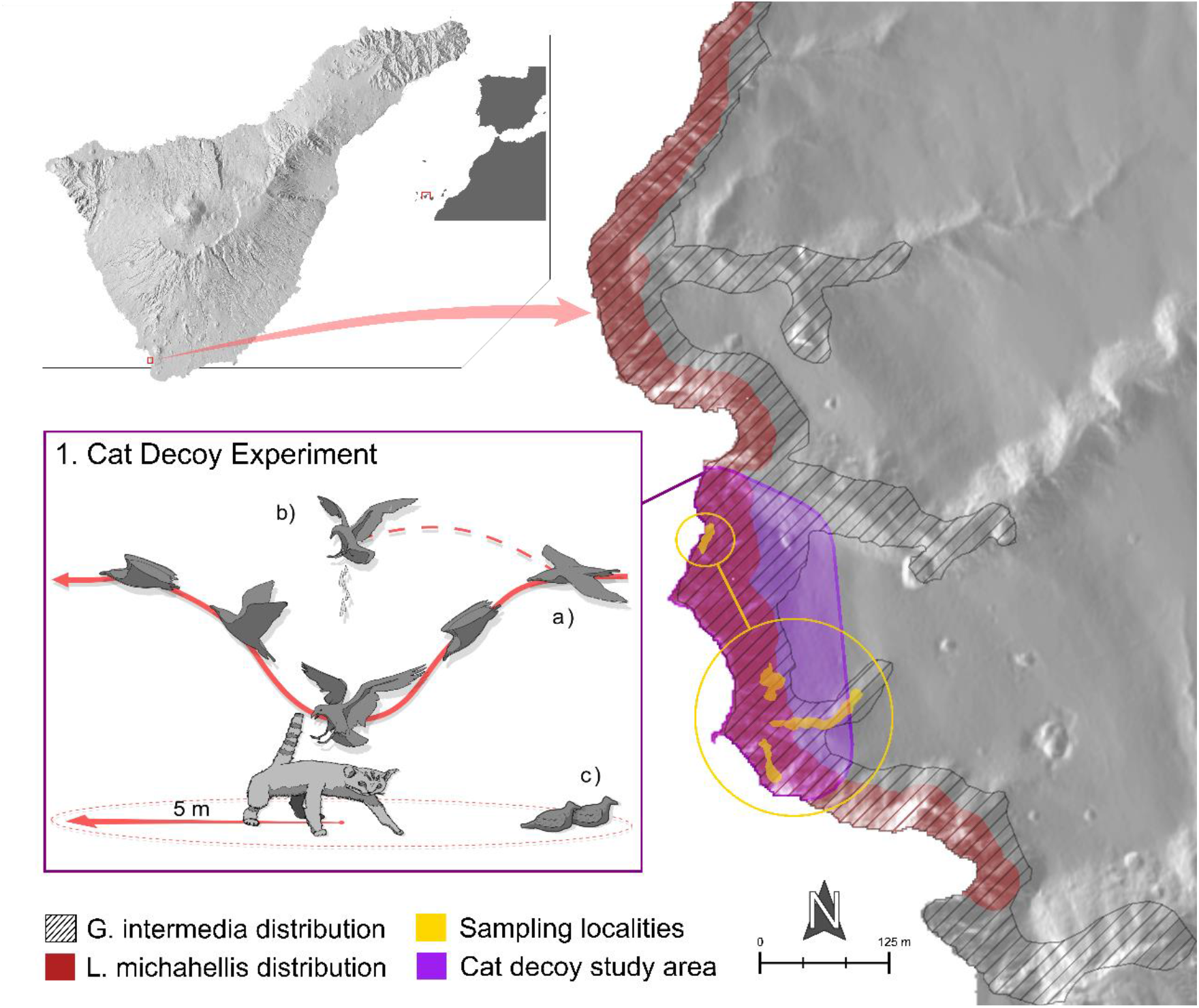
Location and distribution of *Gallotia intermedia* in the Natural Reserve of Guaza Mountain. The red-shaded area describes the distribution of the breeding colony of *Larus michahellis* estimated using on-site observations and high-definition satellite multispectral images (see Supplementary materials S2 for more details). Gridded area marks the described distribution of *G. intermedia* according to Ginoves et al., (2005). Yellow and purple shaded areas mark the sampling localities for isotopes as well as the area used to carry out the anti-predator behaviour experiments. **Panel 1** describes in detail the different types of behaviour considered as aggressions (**a**=dives and **b**= fecal attacks) and the distances from the decoy at which seagulls’ chicks were counted (**c**).

To evaluate our hypothesis, we conducted two distinct yet complementary experiments. To assess whether the populations of *G. intermedia* residing along the coastal cliffs of Guaza are trophically reliant on marine subsidies from the yellow-legged gull colony and the vegetation situated on the platforms and ravines both under and beyond the influence of the seagulls, we conducted a stable isotope analysis (SIA). In a second experiment, we tested whether the colony of *L. michahellis* was effective at keeping feral cats out of this species’ breeding area. To test the anti-predator behaviour of *L. michahellis* towards feral cats, we placed a cat decoy at random locations inside the breeding area of *L. michahellis* and recorded the behaviour of the animals. Finally, based on our results, as well as on the characteristics and conservation status of *G. intermedia* and the rest of the denoted, giant lizards of the Canary Islands, we propose these species to be considered refugee species.

## 2. Material and methods

### 2.1. Study system

Nowadays, the Tenerife speckled lizard naturally occurs in two different populations 30 km from each other, one in the Teno Massif and another one in Guaza Mountain (J.-C. Rando et al., 2004). In this study, we focused on the latter. The coastal cliffs of Guaza Mountain, located on the southern-west side of Tenerife, are part of an eroded volcanic dome which is considered a natural monument protected by the Canarian autonomous government and is included in the European Natura 2000 Network due to its unique geological features and the presence of protected birds’ species under the European Birds Directive (Directive 2009/147/EC) (Figure 1). This volcanic dome is located between two of the biggest touristic localities of the island, Las Americas and El Palm-Mar. At a large scale, the vegetation is mainly formed by a coastal shrubland with a plant association described as *Ceropegio fuscae-Euphorbietum balsamiferae salsoletosum divaricatae*, dominated by *Euphorbia balsamifera* (del Arco Aguilar et al., 2010). Guaza Mountain is located in the arid infra-mediterranean bioclimatic region (del Arco Aguilar et al., 2010), which is characterized by seasonal and scarce precipitation and constant high temperatures. *G. intermedia* inhabits the thin strip of land that contains the coastal cliff area and the transition area between those cliffs and the plateaus (J.-C. Rando et al., 2004; Ginoves et al., 2005) (Figure 1). To obtain an accurate description of the vegetation community in the study area, we conducted a randomized stratified sampling of the vegetation (Elzinga et al., 2001) for each of the topographic features present within the distribution area of the speckled lizard; these features included plateaus, ravines, and cliff platforms (Figure 1, details Supplementary Materials S1). We found differences between the plant communities present at the different topographic features; (Figure S.1); Plateaus were mainly dominated by *Euphorbia balsamifera*, *Limonium pectinatum* and *Frankenia ericifolia* ravines were characterized by *Plocama pendula* and *E. balsamifera* and the cliff platforms were mostly populated by *Patellifolia patellaris* (Table S1-S4).

### 2.3. Sampling

To measure the δ^15^N and δ^13^C composition of *G. intermedia* collected tissue samples of in 4 different localities within its distribution area (2 ravines and 2 platforms) (Figure 1). Reptiles were trapped using pitfall traps of 30x40x50 cm baited with raw tomato and placed using a regular grid of approximately 5 m along the area of the sampling localities. Traps were placed during the early hours of the morning and kept active for 7-8 hours for every trapping session. To avoid animals suffering from excessive heat, we partially cover the entrance of the traps and check them every 30 to 45 minutes, depending on the heat. Each sampling locality was trapped for five consecutive days using 15-23 traps depending on the size of the sampling area. In addition to the pitfalls, we used long rods and lassos to capture the lizards that were more reluctant to fall into the bait traps. Once captured, reptiles were placed in a textile bag and weighted using a spring valance with a range of 300 g and a 0.1 g precision. After this, we recorded some other morphological important metrics such as snout ventral length or SVL (measured ventrally from the tip of the snout to the start of the cloaca), and pilleum width and length using precision callipers and rulers. Tissue samples of *G. intermedia* were collected by cutting, using surgical scissors, a fragment of 2-3 mm from the tip of the tail. *G. intermedia*, as well as the rest of *Gallotia* lizards, use the tail as a fat reservoir as well as a defensive decoy against predators and competitors. For these reasons, tail autotomy is frequent amongst lacertids, and this procedure causes no harm to the individuals (García-Muñoz et al., 2011; Lin & Ji, 2005). Additionally, almost 90% of the captured lizards presented sights of partial or total tail regeneration (Table S5). This sampling method is less invasive than others and allowed us to obtain samples of relatively young tissue from a wide variety of cell types (skin, muscle, bone, etc). Handling of the individuals was carried out in the field, to reduce the interaction time between the animals and the researchers, and under strict sanitary conditions. Once all the samples and metrics were taken, we marked the animals with a temporal identification number (ID in Table S5), using xylene-free nail paint (Boone & Larue, 1999) to avoid re-sampling, and released them in the vicinity of its capture point.

Since we were interested in the role of *L. michaaellis* in shaping the habitat use and diet of *G. intermedia*, we collected tissue samples from yellow-legged seagull nestlings. Tissue samples from chicks were collected by cutting, using a surgical scissor, a small section of the vane from the 2^nd^ and 4^th^ primary feather of each wing (Cherel et al., 2000; Han et al., 2021). We decided to extract the samples from the younglings for various reasons; first, it simplified the sampling and handling of animals (adults of *L. michaaellis* are especially aggressive and cunning, making the direct trapping of the individuals difficult, especially in the harsh terrain of the cliffs, which limits both our movements and the placing of traps); 2) chicks are fast growing individuals that depend fully on their parents for food. Therefore, we expected the isotopic signal of the chick’s feathers to be a strong proxy of the potential marine subsidies that seagulls bring to the breeding area (Cherel et al., 2000). Finally, leaf samples of the dominant plant species in each habitat (platforms, ravines and plateaus) were collected in triplicate (Table S6). In the case of the Cactaceae *O. dillenii* only its fruit was sampled.

### 2.4. Sample processing

Once collected, the tail tips samples of *G. intermedia* as well as plants and seagulls’ feathers samples were rinsed thoroughly in distilled water to remove any surface contaminants before being dried in an oven for 48 h at 45°C, and then ground in a fine powder before the isotopic analyses. Then, 0.3 ± 0.05 mg subsamples of dry powder were weighed in tin capsules for stable isotope analyses. All the tin cups were analysed by combustion at 900°C in a Flash 1112 Elemental analyzer coupled to an isotope ratio mass spectrometer Delta C (ThermoFinnigan, ThermoFisher Scientific) in the lab facilities of the Centres Científics i Tecnològics de la Universitat de Barcelona (www.ccit.ub.edu) in Barcelona, Spain. The abundance of stable isotopes was expressed using the δ notation, where the relative variations of stable isotope ratios are expressed as parts per thousand (‰) deviations from predefined reference scales [atmospheric nitrogen for δ^15^N and Vienna Pee Dee Belemnite (V-PDB) calcium carbonate for δ^13^C]. expressed as δ^y^X = [(R_sample_ – R_standard_) / (R_standard_)], where X is the element, y is the atomic mass of the stable isotope, and R_sample_ is the ratio of heavy to light isotopes (^13^C/^12^C or ^15^N/^14^N) in the sample and R_standard_ is the ratio of the heavy isotope to light isotope in both working reference standards. These analyses were performed with an isotope ratio mass spectrometer. Salt of ammonium sulphate (NH_4_)_2_SO_4_ (IAEAN1, δ^15^N= +0.4‰, and IAEAN2, δ^15^N =+20.3 ‰), L-glutamic acid (USGS40, δ^15^N = -4.5 ‰), and potassium nitrate, KNO_3_ (IAEA-NO-3, δ^15^N = +4.7 ‰) and caffeine (IAEA 600, δ15N = +1.1 ‰. For carbon, secondary isotopic reference materials of known ^13^C/^13^C ratios were used to a precision of 0.2 ‰, and these were namely: polyethylene (IAEACH7, δ^13^C = −32.15‰), sucrose (IAEA CH6, δ^13^C = −10.4‰), L-glutamic acid (USGS40, δ^13^C = −26.4‰), and caffeine (IAEA600, δ^13^C = −27.77‰). These isotopic reference materials were used to recalibrate the system once every 15 samples and were analysed to compensate for any drift over time.

### 2.5. Stable isotopes statistical analysis

Since stable isotopes estimated values turned out to be not normally distributed (Shapiro-wilk test p-value>0.05 in all cases), we used Kruskal-Wallis rank sum test along with a pairwise Wilcoxon rank-sum test to assess the effects of habitat type and *G. intermedia* on the values of δ^15^N and δ^13^C. In addition, the same tests were implemented to test differences in isotopic values among plant samples from different sampling habitats and carbon metabolic pathways (C_3_, C_4_/CAM). A Bayesian stable isotope mixing model was implemented to assess the relative proportional contribution of food resources derived from the three habitats sampled (plateau, platform, and ravine), as well as the contribution of seabird-derived resources, using the package MixSIAR version 3.1.10, (Stock et al., 2018) in R version 3.5.3 (R Core Team, 2021). Both the δ^15^N and δ^13^C values of potential food sources were incorporated into the model, as were the corresponding isotope values of Tenerife speckled lizards. Given that there is no previous information on the diet of *G. intermedia*, we ran the model using uninformative priors that assumed a generalist diet and weighted forage items groups equally (α = 1, 1, 1, 1).

When analysing δ^13^C and δ^15^N in consumer tissue, there is often an offset between the diet and the tissue, termed diet-to-tissue trophic discrimination factor (TDF) or trophic enrichment factor (TEF). These TDFs may vary depending on diet source, growth rates, tissue type, or species (Caut et al., 2008). For accurate diet proportion estimations through the Bayesian isotopic mixing model, it is necessary to use adequate TDFs for each consumer species analysed. Because there are no published trophic discrimination factors for *G. intermedia* or for any other species of the genus *Gallotia*, we used TDFs determined for poikilotherm tissue (δ^13^C 0.4 ± 0.14 ‰ and δ^15^N 2.3 ± 0.2 ‰) according with (McCutchan Jr et al., 2003) and applied in other lizards’ diet studies (Murray et al., 2016).

We estimated the proportion of trophic resources from the three habitats to lizards’ diet, grouping into four forage item groups by the isotopic values of plants by habitat and plant metabolic pathway, since C_3_ or CAM/C_4_ plants differ significantly in their δ^13^C values, as well as the seabird-derived resources. To estimate the proportion of the latter we use as forage item group the isotopic values of seagull chicks by subtracting beforehand the trophic discrimination factor (TDF) calculated on feathers of *L. michahellis* when feeding on marine prey according to Ramos et al. (2009). All the isotopic values of the different forage item groups, as well species their species composition are shown in Table S6. A Markov chain Monte Carlo (MCMC) sampling by running three replicate chains, each with 100,000 iterations, with a burn-in of 50,000 iterations and thin by 50 was used as a setup in MixSIAR. Diagnostic tests (Gelmin– Rubin, Heidelberger–Welch and Geweke) and trace plots were examined for model convergence.

### 2.6. Body-guard effect sampling and analysis

To test the potential role of the yellow-legged seagulls in the protection of *G. intermedia* from predators we placed a cat decoy in 15 different locations along *L. michahellis* breeding colony. For the cat decoy, we used a staffed feral cat fixed to a wooden plank in the stand position (Figure 1 and S3.c). This decoy was placed at random in the breeding colony by one of the members of the team while another researcher was strategically hidden from the seabirds within the direct sighting line to the decoy. Once the decoy was placed, we waited until the yellow-legged seagulls detected it, usually marked by multiple alarm songs by hovering birds, and then recorded the number and type of aggressive interaction against the cat decoy (similar to Gameiro et al., 2022). These interactions were classified into three basic types; 1) dives, in which seagulls perform a rapid “diving” flight towards the decoy and try to hit it with their legs; 2) Faecal attack, in which the birds defecated in mid-flight trying to hit the decoy; and 3) ground attacks, in this instance the animals try to land behind the decoy, out of its potential sight, and directly peck at the tail and rear legs of the decoy (Figure 1). This type of behaviour has been previously described in the literature as aggressive or territorial defensive behaviour (Frixione & Salvadeo, 2021; Larsen, 1991; Minias et al., 2020). To reduce our disturbance in the breeding colony, and to avoid potential behavioural changes in the individuals (Gaynor et al., 2018), we retired the decoy after two minutes of the first aggressive interaction. Similarly, if the decoy was detected but no attack was recorded during the first 3 minutes after its detection, we also retired the decoy. Since we hypothesized that the anti-predator behaviour is greater within the breeding area of *L. michahellis*, once the decoy was hidden, we counted the number of young chicks located in a radius of 5 m from where the decoy was placed.

Time was controlled using digital chronometers and attacks were recorded using a combination of binoculars and hidden wide-angle video cameras located in the vicinity of the decoy (GoPro Hero3 model CHDHN-301). The feral cat decoy used for this experiment presented the usual stripped brownish colouration common amongst wild-born feral cats (Figure S3.c). This animal was accidentally run over and killed on the island of Gran Canaria in 1985. It was later donated and prepared for the collection of the Zoology Department of the University of La Laguna, where it was returned after the experiment.

To test whether the number, type, and probability of aggressions towards the decoy was related to the density of birds within the breeding colony, we fitted a generalized linear models (GLMs) with a Poison and Binomial distributions using counts of the different attacks (Dives; Faecal-attack; ground attack; and total number of aggressions) and it’s binomial form (1= attack, 0= no attack) as response variables, using the number of yellow-legged chicks as the explanatory variable. Seagull nests, and therefore chicks, are not uniformly distributed along the breeding colony. Terrain features such as slopes can determine where nests and animals are located (Momberg et al., 2023) (see supplementary materials S2). Due to this, we added terrain ruggedness, calculated as the mean of the absolute elevation difference between a cell and its 8 surrounding neighbouring cells (Wilson et al., 2007), into the models as a co-variable.

All models were fitted by maximum likelihood, using an adaptative Gauss-Hermite quadrature approximation, implemented in the *lme4* R package version 1.1-2368 (Bates et al., 2015). To satisfy the assumptions of linear relationships of the residuals and to reconcile the different scales of the variables, terrain ruggedness was standardized by subtracting the mean and dividing by the standard deviation. Model performance was measured using Nagelkerke’s r^2^ (Nagelkerke & others, 1991) implemented in the R package *performance* version 0.9.1 (Lüdecke et al., 2021). The statistical significance of the model effects was calculated using the Type II Wald Chi-square test, implemented in the *car* R package version 3.1-0 (Fox & Weisberg, 2019). All analysis, data manipulation, and spatial pre-processing were carried out using the R statistical software version 4.3.1 (R Core Team, 2021).

## 3. Results

### 3.1. Isotopic characterization of the Tenerife speckled lizard, habitat, and trophic resources

In total, we analysed the δ^15^N and δ^13^C of 79 plant samples from 12 different species from 10 families, 10 feather samples from gull chicks (*L. michahellis*) and a total of 28 samples of Tenerife speckled lizards (*G. intermedia*) from Guaza cliffs. Lizards’ samples were archived between July and September from two different habitats, (n=7) from isolated platforms, and (n=21) samples from ravines. Values of δ^15^N for *G. intermedia* ranged from 11.53‰ to 15.94‰ and values of δ^13^C ranged from -21.55‰ to -18.41‰, and their mean values expressed both as (mean ±SD) were 14.03 ± 1.01‰ for δ^15^N and -19.76 ± 0.9‰ for δ^13^C, (Figure 2.a). Stable isotope ratios of δ^15^N from platforms lizards ranged widely from 11.93‰ to 14.81‰ (mean ± SD = 13.5 ± 0.99 ‰) and from 11.53‰ to 15.95‰ (mean ±SD = 14.24 ± 0.97 ‰) in ravines lizards, while δ^13^C values from platforms specimens ranged from -20.45‰ to -18.41‰ (mean ± SD = -19.1 ± 0.83‰) and from -21.55‰ to -18.78‰ (mean ± SD = -20.01 ± 0.83 ‰), for ravine habitat specimens. Lizards inhabiting ravines showed slightly higher values of δ^13^C those of platforms (Kruskal-Wallis χ^2^= 5.3265, df=1, p=0.021). There were no statistically significant differences in the values of δ^15^N between reptiles inhabiting different habitats (χ^2^= 2.61, df=1, p=0.1056).

**Figure 2.**
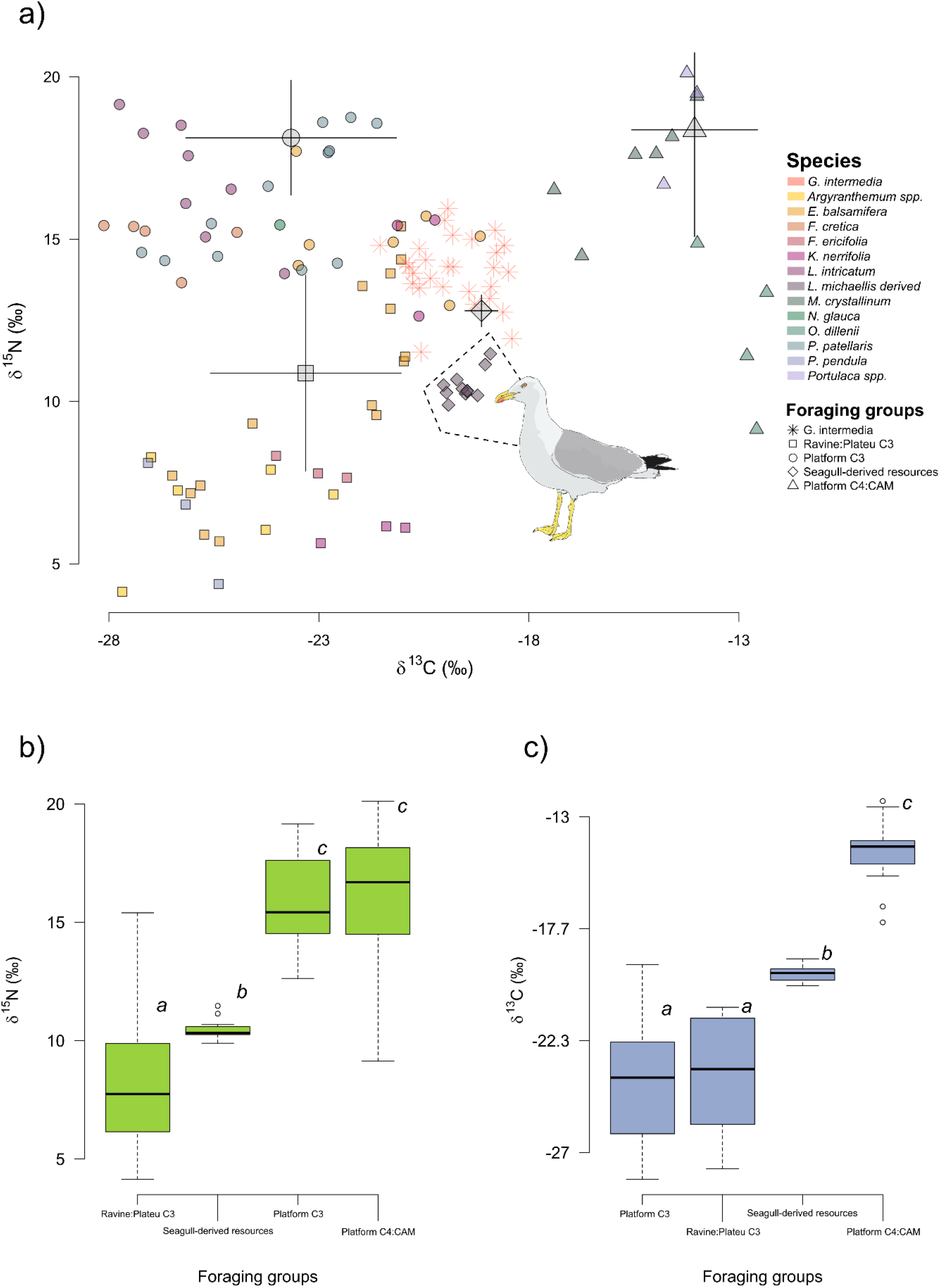
Values of δ^15^N and δ^13^C for each species (**a**) and habitat sampled (**b-c**) during the study of the diet of *Gallotia intermedia*. Points on panel **a**, represent the raw values obtained from the tissue samples of *G. intermedia* and the rest of the potential trophic resources. Species are differentiated by colours whereas the distinct geometrical shapes represent the different foraging groups. The mean ± the standard deviation δ^15^N and δ^13^C calculated by the MixSiar analysis taking into consideration the Trophic Discrimination Factors (oversized grey shapes and lines of each foraging group) are presented along with the raw δ^15^N and δ^13^C for the different species. Panel b and c represent the total values of δ^15^N and δ^13^C for the 4 types of Foraging groups included in the analysis. Lowercase letters along the boxplot represent pairwise statistically significant groups.

Vegetation samples were classified as either C_3_ (δ^13^C values within the −22‰ to −30‰ range) or C_4_/CAM (δ^13^C values in the −10‰ to −14‰ range). In this manner, δ^13^C values of individual C3 plant species ranged from −28.12‰ to −19.16‰ and from −17.40‰ to −12.33‰ for C_4_/CAM plants. δ^15^N values for C3 plants ranged from 4.14‰ to 19.15‰ and from 9.14‰ to 20.11‰ for C_4_/CAM plants (Table S7). Seagull chick feather samples ranged in δ^13^C values from -18.92 ‰ to -20.03‰ and from 9.89 to 11.47 for δ^15^N values (Table S7).

The composition of δ^15^N and δ^13^C across the four foraging items groups formed by plants metabolic pathway and habitat were statistically significant different in both isotopes ratios values (δ^15^N χ^2^=59.269, df=3, p<0.001; and δ^13^C χ^2^= 52.315, df=3, p<0.001) (Figure 2.a-c). Pairwise comparisons revealed statistically significant differences among δ^15^N values in platforms C_3_ and C_4_/CAM plants and ravine-plateau plants, as well as from seagull-derived resources. Platform plants showed in general higher δ^15^N values (15.89 ± 2.20‰) than ravine/plateau plants (8.57 ± 4.25‰), even than seagull-derived resources (10.49 ± 0.45‰) (Table S7). Whereas for δ^13^C values only C_3_ plants showed no significant differences (p>0.05), independently of their habitat, being statistically different from C_4_/CAM plants (-14.45 ± 1.49‰) which presented the highest values among plants followed by seagulls-derived resources (-19.53 ± 0.37‰) (p<0.001), see Table S7 for full numerical values of pairwise Wilcoxon rank-sum test. The MixSIAR model estimated that the mean (±SD) contribution of seabird-derived resources dominated the dietary composition of the Guaza mountain population of *Gallotia intermedia* (62.2 ± 14.3%), following by C_3_ plants of platforms (15 ± 0.42%) and C_3_ plant from ravines and plateau habitats (12 ± 10.1%) (Table 2, Figure. 2.a).

**Table 1:** Mean (± standard deviation) values (δ^15^N and δ^13^C) of foraging item groups used in the mixing model MixSIAR to calculate diet contributions of *Gallotia intermedia*. n: number of samples analysed. Group 1: *E. balsamifera*, *F. cretica*, *K. neriifolia*, *L. intricatum*, *N. glauca*, *P. patellaris*. Group 2: *M. crystallinum*, *O. dillenii*, *Portulaca* spp. Group 3: *Argyranthemum* spp., *E. balsamifera*, *F. ericifolia*, *K. neriifolia*, *P. pendula*. Group 4: *L. michahellis* chicks’ values after TDF subtraction.

| Forage item group | Mean ± $\delta^{13}\text{C}\text{‰}$ | Mean ± $\delta^{15}\text{N}\text{‰}$ | n |
| --- | --- | --- | --- |
| Group 1: Platform C<sub>3</sub> | -24.06 ± 2.51 | 15.82 ± 1.75 | 36 |
| Group 2: Platform C<sub>4</sub>/CAM | -14.46 ± 1.49 | 16.07 ± 3.29 | 13 |
| Group 3: Plateau/Ravine C<sub>3</sub> | -23.71 ± 2.27 | 8.57 ± 3.01 | 30 |
| Group 4: Seabirds-derived resources | -19.5 ± 0.37 | 10.5 ± 0.45 | 10 |

**Table 2:** Proportional contribution of each forage item group to the diet of *Gallotia intermedia* estimated using MixSIAR models. Mean ± SD, 50% quantiles (median) and 95% Bayesian credible intervals (CI).

| Forage item group | Mean ± SD | 50% | 95% CI |
| --- | --- | --- | --- |
| Group 1: Platform C<sub>3</sub> | 15 ± 0.42 | 15.2 | 5.8-22.7 |
| Group 2: Platform C<sub>4</sub>/CAM | 10.9 ± 6.6 | 9.6 | 1.6-27.6 |
| Group 3: Plateau/Ravine C<sub>3</sub> | 12 ± 10.1 | 9.2 | 0.06-38.7 |
| Group 4: Seabirds-derived resources | 62.2 ± 14.3 | 65.6 | 24.8-80.5 |

### 3.2. Bodyguard effect

We recorded a total of 165 aggressive interactions between *L. michahellis* and the feral cat decoy across 14 different locations within the breeding colony; 135 dives; 27 faecal attacks; and 3 ground or direct attacks (the later ones were not included in the analysis due to its low frequency). The frequency of dives (Poisson family models), as well as the total number of attacks, were higher in the areas with a higher number of *L. michahellis* chicks (Type II Wald Chi-square test; χ2 = 77.130, d.f = 1, P < 0.001; and χ 2 = 64.397, d.f = 1, P < 0.001, respectively) (Figures 3.a and 4.a). Model predictions for the frequency of dives and total number of attacks showed that these are mainly restricted to the limits of the yellow-legged seagull breeding colony, being strongest in the core areas of the breeding colony and fading away towards their edges (Figures 3.c and 4.c). However, Fecal attacks were independent of the number of chicks (χ2 =0.073, d.f = 1, P=0.785), and the ruggedness of the terrain (χ2 =1.027, d.f = 1, P=0.31). Model predictions showed that faecal attacks were restricted to *L. michahellis* breeding area but randomly distributed within its limits.

**Figure 3.**
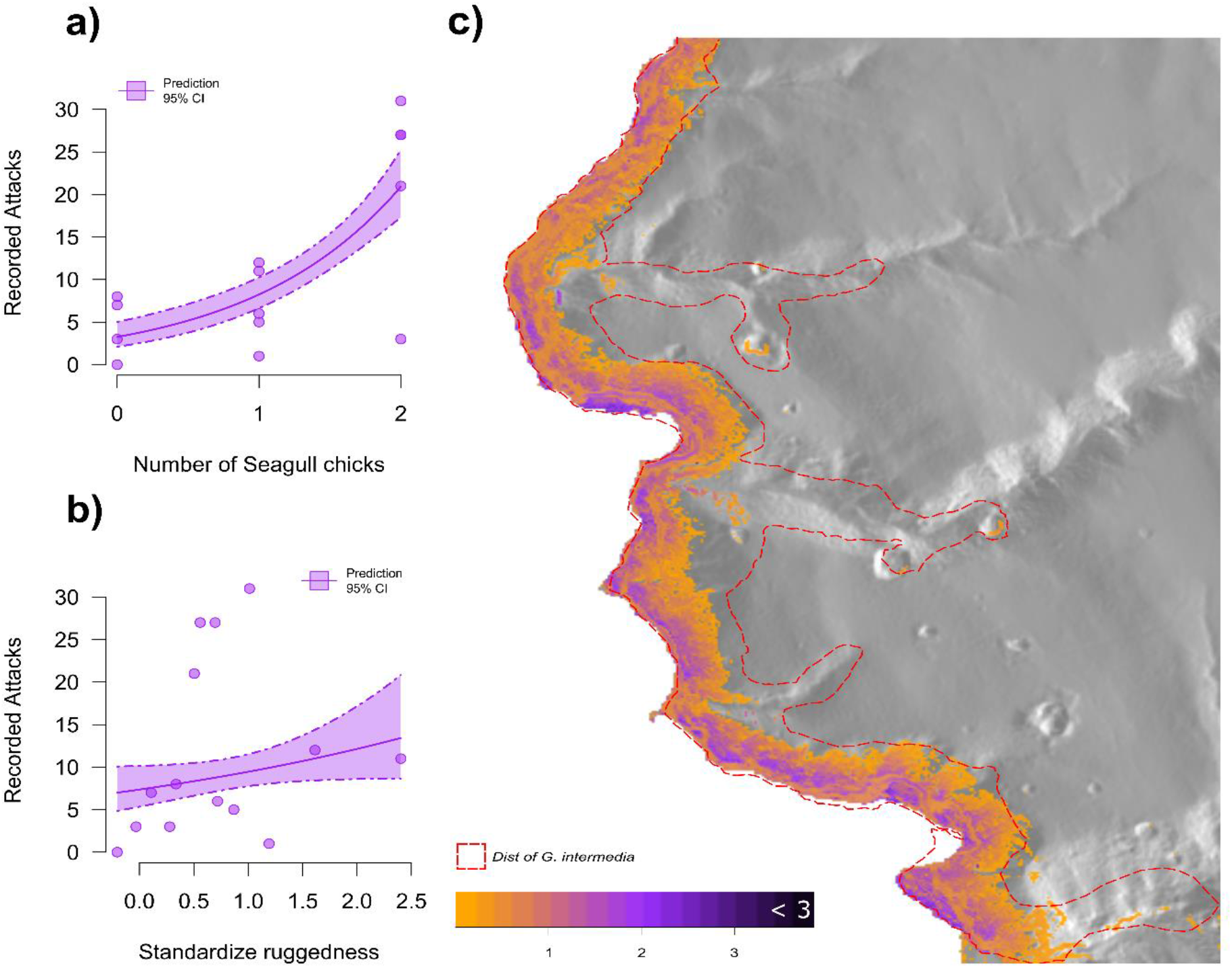
Results of the anti-predator behaviour analysis. Partial responses of the Poisson generalized linear models describing the relationship between the number of seagull chicks (**a**) and standardised topographic ruggedness (**b**) with the total number of recorded attacks (Dives + Fecal attacks). Data points are presented along with predicted responses and 95% confidence intervals. Spatial predictions of the number of *Larus michahellis* attacks (**c**), dotted red lines represent the approximate distribution of *G. intermedia* according to Ginoves et al., (2005).

**Figure 4.**
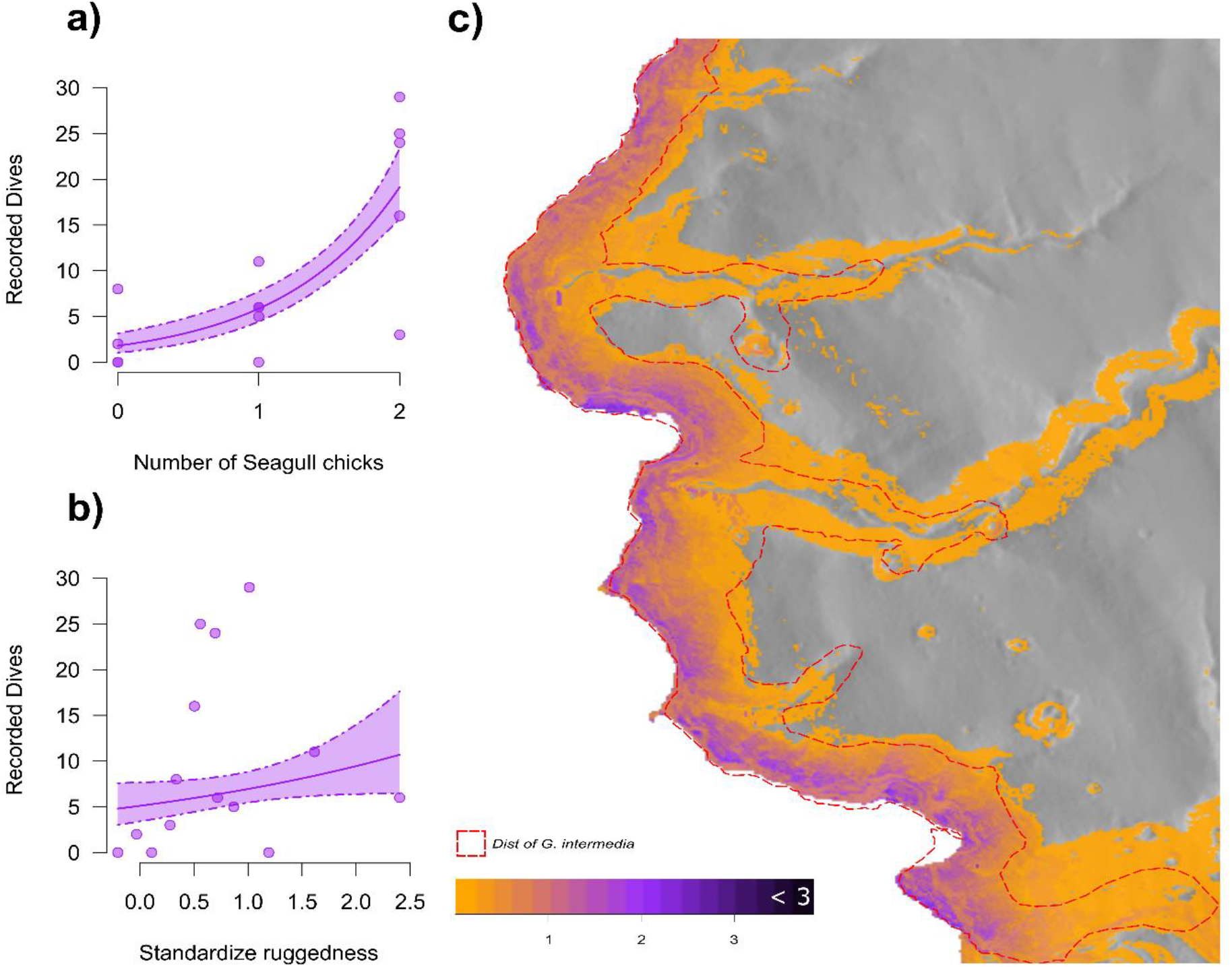
Results of the anti-predator behaviour analysis. Partial responses of the Poisson generalized linear models describing the relationship between the number of seagull chicks (**a**) and standardised topographic ruggedness (**b**) with the number of recorded dive attacks. Data points are presented along with predicted responses and 95% confidence intervals. Spatial predictions of the number of *Larus michahellis* attacks (**c**), dotted red lines represent the approximate distribution of G. intermedia according to Ginoves et al., (2005).

In terms of *L. michahellis* attack probability (Binomial family models), models predicted an independent distribution of the probability of an attack taking place (dives and fecal attacks) in relation to the number of seagull chicks (χ2 =0, d.f =1, P=0.999). The same pattern was observed for dives and fecal attacks separately (χ2 =2.307, d.f =1, P=0.128 and χ2 =0.17, d.f =1, P=0.676 respectively).

## 4. Discussion

In this study, we showed how direct and indirect ecological interactions allow a critically endangered reptile to inhabit and maintain relatively stable populations in a suboptimal habitat. Through two different experiments, a dietary stable isotopes analysis, and an ecological behaviour study, we showed how the Tenerife speckled lizards interact with a yellow-legged seagull colony to obtain food, and protection against alien predator species. The stable isotope analysis showed that revealed that *G. intermedia* was mainly feeding in the area under the direct influence of seagulls’ guano and taking advantage of the seagull-derived food resources that they provide directly or indirectly. Additionally, the behaviour experiment showed that *L. michahellis* displayed an aggressive antipredator behaviour against predators within the limits of its breeding area with an increase in the frequency of aggressive behaviours correlated with the abundance of seagull’s chicks.

The effects of seabird-derived resources on lizard populations have been found to positively affect different parameters such as reproduction, body size, density and intraspecific antagonistic behaviour (Larsen, 1991; Pafilis et al., 2009, 2011; K. M. Richardson et al., 2019; Stadler et al., 2023). The patterns of stable nitrogen and carbon isotopes found on *G. intermedia* back to those past results. Furthermore, the patterns of stable carbon and nitrogen found on the different trophic resources as well as on *G. intermedia,* showed us how the latter is using its environment (Cherel et al., 2000; Grundler et al., 2017; Han et al., 2021). The vegetation in areas under the direct influence of *L. michahellis* colony shows a completely different isotopic signal. The high values of δ^15^N of the plants located nearby or directly inside the colony show a clear influence of nitrogen derived from guano, which exhibited δ^15^N values ∼7‰ higher than those on the areas out of seagull colony (Figure 2.a-b, Table S.7). The vegetation from platforms were as expected showed the highest δ^15^N values, even higher than those of the seagull chicks (mainly piscivores) and lizards. It is well known that seabird colonies, through the deposition of guano, rich in nitrogen and phosphorous compounds, induce profound changes in soil properties, including eutrophication, salinization, acidification and nutrient imbalances, modifying the plant communities that live there (Anderson & Polis, 1999; Ellis, 2005; Wainright et al., 1998; Wait et al., 2005). As a consequence, plants enriched in δ^15^N improve their quality and quantity and significantly increase the δ^15^N content in all components of the food webs of coastal and island ecosystems through multi-trophic cascading effects (Barrett et al., 2005; Fukami et al., 2006). In the case of *G. intermedia*, its diet seems to depend on approximately 25% of plants that live in the area influenced by the seagull colony. This limited contribution may be indicative of the sparse vegetation cover in these areas, where C_4_/CAM plants such as *Portulaca* spp., *M. crystallynum* and *O. dillenii* predominate, along with C_3_ plants such as *P. patelleris*, *L. itrincatum* and *K. neriifolia*. Many of these plants have an annual cycle, appearing only during the wet season. In contrast, the prickly pear *O. dillenii* (cactacea) produces ripe fruits nearly year-round, which are consumed by *G. intermedia* (pers. obs.). However, the distribution of *O. dillenii* on the Guaza cliff platforms is limited to a few locations. Conversely, the prickly pear is a non-native invasive species in the Canary Islands, leading to calls for its control and eradication in protected areas (García-Gallo et al., 2009). Given our findings, the eradication of this species in areas inhabited by the Tenerife speckled lizard could potentially jeopardize its survival. Therefore, we recommend considering the trophic and other ecological relationships that may exist between naturalized invasive plant species and threatened endemic species before implementing eradication plans.

The MixSIAR model suggests that the diet of *G. intermedia* is primarily comprised of food items derived from seagulls (∼62%). These resources displayed δ^13^C levels of approximately -18‰, which are markedly different from the δ^13^C levels of C3 plants (∼ -24‰) and C4/CAM plants (∼ -14‰). These levels indicate that the carbon is sourced from marine and epipelagic food sources in the area, such as epipelagic fishes like *Trachurus picturatus* and *Scomber colias*, which have δ^13^C values ranging between – 19‰ and -20‰ (Romero et al., 2021). Similarly, the δ^13^C values of *G. intermedia* (∼ – 20 ‰) are indicative of this marine epipelagic δ^13^C signature. This suggests that most of the foraging activity of *G. intermedia* is taking place in the areas under the direct influence of the yellow-legged seagulls, despite the majority of speckled lizards being caught in ravines, where vegetation cover and diversity can offer a more diverse range of foods resources. Seagull-derived resources may encompass a wide range of foods, including remains brought directly by the parents to the chicks (such as epipelagic fish and cephalopods), which may be regurgitated and left outside the nest, as well as remains of chick’s or adult seagulls carcasses, eggshells, seagull droppings and scavenger insects that feed on these remains and move out of the colony area dispersing this marine subsidy through the food chain. Future dietary studies with trophic markers such as those used here can provide a greater resolution if more effort is made to sample all possible food sources used by this species throughout the year, obtain food consumption data derived from fecal analysis (priors), and generate species-specific or genus-specific TDFs to improve the fit of Bayesian isotopic mixing models.

We cannot discount the possibility that ravine areas serve as dispersal zones for juvenile lizards originating from the cliff. Ontogenetic habitats shift in lizards, where juveniles actively avoid the presence of adults due to competition, cannibalism or select other habitats according to their different life trait requirements have been documented in multiple lizard species (Delaney & Warner, 2017; Jenssen et al., 1989; McMann & Paterson, 2003; San-Jose et al., 2016; Stamps, 1983). In the context of *G. intermedia*, the forced removal of smaller individuals from cliff regions, where they benefit from protection and food supplied by gulls, could heighten their susceptibility to predation and diminish the fitness of these juveniles in ravines. This may ultimately lead to these ravines becoming sinks for the population. Unfortunately, the low number of captures in our study does not allow us to investigate this question, so future research should be carried out to further investigate the possible habitat selection of *G. intermedia* by age and sex and the structuring of their populations.

Additionally, the results from the anti-predator behaviour experiment further back the potential positive interaction between the breeding colony of *L. michahellis* and *G. intermedia*. Although the probability of attack was randomly distributed within the colony, the frequency of attacks was strongly correlated with the abundance of the seagull’s chicks. Similar mobbing behaviours have been recorded in other seabirds’ colonies (Clode et al., 2000; Larsen, 1991), and in some cases, it has been reported that other species use this indirect protection to their advantage (Gameiro et al., 2022; Guidos et al., 2023). For example, Monk parakeets (*Myiopsitta monachus*), an IAS built their nest under those of white storks (*Ciconia ciconia*) which indirectly offers protection against predators (Hernández-Brito et al., 2020). The anti-predator behaviour of *L. michahellis* wasn’t limited only to our cat decoy. During the months of fieldwork, we observed aggressive behaviour towards a wide range of species, from kestrels to Eleonora falcons, egrets, and biologists (see Supplementary Materials Figure S3.a-c and Figures S4). Again, these types of interactions can help us explain why *G. intermedia*, a critically endangered island species, can maintain its population in an area under feral cats’ pressure and surrounded by huge touristic developments. The scale and reach of the effects of feral and domestic cats on the populations of *G. intermedia* are still unknown, however, the limited number of studies on the subject have shown that the Tenerife speckled lizard is a relatively common prey for the cats that roam the coastal areas of Guaza Mountain (Albaladejo et al., 2015; Flores Ravelo & Rando Reyes, 2021). Although *L. michahellis* can help to mitigate the damage, at least during its breeding season, previous monitoring of *G. intermedia* populations in the protected area of Guaza Mountain has shown a potential population decline of more than 32% over the past 18 years (Albaladejo et al., 2015). Therefore, active conservation actions that take into consideration the complex interactions between the species are fundamental for pushing endangered species away from extinction.

Although we focused just on one species and a set of small populations, our findings are of interest for the conservation and study of other critically endangered lizard species that share similar characteristics and requirements with *G. intermedia*. This is the case, for example, of *Gallotia bravoana* (Mateo et al., 2009a) and *Gallotia simonyi* (Mateo et al., 2009b). These endemisms of La Gomera and El Hierro islands, respectively, are also severely affected by feral cats (Afonso & Mateo, 2009; García Márquez et al., 1997) and their natural populations are restricted to isolated and harsh cliffs (Afonso & Mateo, 2009; Albaladejo et al., 2015; Rica, 1981) occupied by different breeding populations of seabirds (e.g., marsh and shearwaters, seagulls). Understanding which were the conditions that allowed this species to maintain wild populations in such harsh conditions can aid the reintroduction success of future populations by not only looking for the right geographic conditions (isolation and difficult access to invasive species) but also for adequate biotic interactions (Bogoni et al., 2019; Renzi et al., 2019). Furthermore, populations of the giant lizard of Gran Canaria (*Gallotia stehlini*) are being decimated by the introduction of the California Kingsnake (*Lampropeltis californiae*) (Piquet & López-Darias, 2021) and so it is likely that shortly this species also will face the same challenges that the rest of endemic giant reptiles of the Canary Islands. In this context, refugees can be identified and preserved in advance, and individuals can be introduced in these suboptimal habitats to preserve the genetic diversity of the species (Braidwood et al., 2018) while other conservation actions are undergoing to try to halt the expansion of the California Kingsnake to the rest of the island (Piquet et al., 2021).

At a global scale, habitat destruction and invasive alien species are one of the main causes of biodiversity loss (Dirzo et al., 2014; WWF, 2020). In this scenario, we should expect more species to be pushed into suboptimal or relict habitats. As we have shown with this work, understanding which elements allow species to survive in such conditions is of paramount importance for the correct planning and management of conservation actions. Therefore, a focus on disentangling the hidden biotic and abiotic interactions that allow species to inhabit these suboptimal habitats can help us to better prioritize habitat conservation, reintroduction areas, and long-term species management. For this, it is fundamental that basic ecological studies are conducted alongside biodiversity conservation (Brussard, 1991; Fontúrbel et al., 2018).

Overall, we found that populations of critically endangered species inhabiting a relict or suboptimal habitat rely on indirect ecological interactions to maintain their populations. Our results suggest not only an important interconnectivity between the yellow-legged seagull colony and the Tenerife Speckled lizard but also that the realised niche of *G. intermedia* is defined by this interaction. Our observations further back the importance of basic ecological research to understand the mechanism behind the distribution and survival of endangered species, and the importance of this knowledge for the correct application of conservation actions. The results of this study can further help understand the patterns of distribution of other endemic and endangered reptiles of the Canary Islands as well as to design long-term conservation strategies. At a global, since we expect more species to be pushed to suboptimal habitats, we suggest that hidden biotic interactions such as the one we describe here might be of paramount importance for the conservation of biodiversity and the planning of conservation actions.

## Declaration of Competing Interest

The authors declare that they have no known competing financial interests or personal relationships that could have appeared to influence the work reported in this paper.

## Acknowledgements

We thank all the people who helped us during the fieldwork (M. Gil-Velasco, J. Marrero, N. Álvarez-Quintero). Also, to Drs. D. Hernández-Teixenor and Miquel Lombarte for their help with the SIA analyses. To Dr. A. Martín who provided the essential logistical support that made this research possible, as well as lending us the decoy cat “Fermín” for the experiments. We would also like to thank the staff of As— Tonina for the management of the project.

## Funding

This research has been funded by The Mohamed bin Zayed Species Conservation Fund within the 2016 call, project number 152511662. A. E. has been funded by the Actions of the Ministry of Universities under the application 33.50.460A.752 and by the European Union Next Generation EU/PRTR through a Margarita Salas postdoctoral contract of the University of Vigo.

## Author contributions

G. A. R.: conceptualization, formal analysis, investigation, methodology, software, writing the original draft, writing the review and editing A. E.: conceptualization, formal analysis, funding acquisition, investigation, methodology, project administration, writing the original draft, review and editing. A. V. P. C.: formal analysis, investigation.

## Data Availability

Data will be made available on request.

## Supplementary materials *Gallotia intermedia*

### S1. Characterization of the vegetation in the habitat of *G. intermedia*

The vegetation study was conducted during two sampling sessions at the end of the dry season in early March 2016. Stratified random sampling of vegetation was carried out for each of the considered topological features found within the potential distribution area of *G. Intermedia* (Fig 1 main text); cliff platforms; ravines; and plateau. We carried out this sampling by placing five-line intercept transects of five meters long each. The initial position as well as the direction of the transects was randomly selected by throwing a 30 cm wooden ruler into the air. The landing point was set as the starting point of the transect whereas the orientation of the ruler marked the direction of the transect. Vegetation cover was measured along each line intercept transect by noting the point along the tape where the canopy of each plant species begins and the point at which it ends. The percentage of cover for each plant species in the line transect was then calculated as the proportion of transect covered by each plant species (Elzinga et al., 2001).

To characterize and detect potential differences in plant communities between different topological features, we applied a multivariate analysis approach. Species cover data was log (x+1) transformed and used to construct a similarity matrix using the Bray-Curtis dissimilarity index. The resulting matrix was then employed to perform cluster and non-metric multidimensional scaling (nMDS) analyses. Variation in plant community composition between topological features was subsequently tested using a one-way analysis of similarities (ANOSIM). Similarities between plant communities were measured using the ANOSIM R statistic, calculated using 999 Monte Carlo permutations. The R-value returned by this test represents the degree of dissimilarity between the plant communities found in the topographic features. ANOSIM R values close to “1” suggest dissimilarity between groups while an R-value close to “0” suggests an even distribution of high and low ranks within and between groups (Clarke, 1993). In addition, to explore the relative contribution of plant species to the characterization of communities, we applied a similarity percentage analysis (SIMPER). This analysis tests which species contribute to the difference between communities and how much of that difference can be explained by each species. All these analyses were carried out using the R package *Vegan* version 2.6-4 (Oksanen et al., 2022).

#### Results

The results from the nMDS, as well as the hierarchical clustering of the Bray-Curtis dissimilarity matrix, suggest that the topography of the environment is a key element in the delimitation and configuration of plant communities along the distribution area of *G. intermedia* in the Natural Reserve of Guaza Mountain (Figure S1 and Table S1-4). This was further confirmed by the results of the ANOSIM analysis which confirmed a strong difference between plant communities according to the topographic features (R-value=0.87; p-value=0.001; with 999 permutations). The SIMPER analysis showed that *Patellifolia patellaris*, *Euphorbia balsamifera*, and *Argyranthemum foeniculaceum* were the main contributors to differentiate the platform from the plateaus and ravines plant communities (Table S1). In the case of plateaus and ravines, *Arguranthemum foeniculaceum*, *Neochamaelea pulvurulenta*, and *Plocama pendula* were the species contributing to the differentiation between those plant communities (Table S1).

**Table S1.** Aggregated abundances for the plant species were detected during the stratified sampling of the 4 different localities. The most abundant species for each locality are marked in bold, and the total vegetation cover for each locality is shown at the bottom of the table.

|  | Ravine | Plateau | Platforms |  |
| --- | --- | --- | --- | --- |
| **Species** | **BNAME** | **CNAME** | **NAME** | **NICOTIANA** |
| *Aizoon canariense* | 0 | 0 | 0.0056 | 0 |
| *Argyranthemum foeniculaceum* | 0.186 | 0.0032 | 0 | 0.012 |
| *Astydamia latifolia* | 0 | 0 | 0 | 0.024 |
| *Aeonium sp* | 0.008 | 0 | 0 | 0 |
| *Cenchrus ciliaris* | 0.04 | 0 | 0.02 | 0.024 |
| *Euphorbia balsamifera* | 0.184 | **0.3288** | 0.174 | 0.084 |
| *Fagonia cretica* | 0 | 0.004 | 0.01 | 0 |
| *Frankenia ericifolia* | 0 | 0.0908 | 0 | 0 |
| *Kleinia nerifolia* | 0.044 | 0.06 | 0.022 | 0 |
| *Limonium pectinatum* | 0.008 | 0.144 | 0 | 0 |
| *Neochamaelea pulvurulenta* | 0.112 | 0 | 0 | 0 |
| *Opuntia dillenii* | 0.0528 | 0.096 | 0 | 0 |
| *Patellifolia patellaris* | 0 | 0 | **0.2148** | **0.586** |
| *Plocama pendula* | **0.2** | 0 | 0 | 0 |
| *Astydamia latifolia* | 0 | 0 | 0.0092 | 0 |
| *Zygophyllum fontanesii* | 0 | 0 | 0.0572 | 0 |
| *Reseda scoparia* | 0 | 0.02 | 0 | 0 |
| *Schizogyne sericea* | 0.008 | 0.056 | 0 | 0 |
| *Volutaria canariensis* | 0.0016 | 0 | 0 | 0 |
| Total vegetal cover | 0.8444 | 0.8028 | 0.5128 | 0.73 |

**Table S2.** ANOSIM results for the plant communities inhabiting **platforms** and **plateau** habitats along the distribution area of *G. intermedia*. The average contribution of species to the differentiation between plateaus and platforms plant communities (**Average**), is presented along with the standard deviation of these contributions (**sd**); the relative abundance of species (**Abund**); the cumulative contribution of species (**c.sum**); and the p-values (***p***) of the ANOSIM permutations, in this case 999. Species with no abundance or contribution to the communities have been removed from the table.

| **Species** | **Average** | **sd** | **Abund** | **c.sum** | **p** |
| --- | --- | --- | --- | --- | --- |
| *Patellifolia patellaris* | 0.183 | 0.052 | 5.008 | 0.209 | 0.001 |
| *Euphorbia balsamifera* | 0.128 | 0.094 | 1.949 | 0.355 | 0.017 |
| *Limonium pectinatum* | 0.101 | 0.087 | 0.000 | 0.471 | 0.004 |
| *Frankenia ericifolia* | 0.092 | 0.081 | 0.000 | 0.576 | 0.001 |
| *Opuntia dillenii* | 0.068 | 0.086 | 0.000 | 0.654 | 0.041 |
| *Schizogyne sericea* | 0.055 | 0.071 | 0.000 | 0.717 | 0.011 |
| *Reseda scoparia* | 0.049 | 0.062 | 0.000 | 0.772 | 0.001 |
| *Mesembryanthemum nodiflorum* | 0.042 | 0.057 | 1.288 | 0.820 | 0.496 |
| *Kleinia nerifolia* | 0.040 | 0.069 | 0.403 | 0.866 | 0.421 |
| *Argyranthemum foeniculaceum* | 0.027 | 0.045 | 0.343 | 0.896 | 1.000 |
| *Cenchrus ciliaris* | 0.025 | 0.052 | 0.804 | 0.926 | 0.895 |
| *Fagonia cretica* | 0.025 | 0.042 | 0.326 | 0.954 | 0.187 |
| *Astydamia latifolia* | 0.021 | 0.044 | 0.676 | 0.979 | 0.576 |
| *Zygophyllum fontanesii* | 0.011 | 0.033 | 0.318 | 0.991 | 0.677 |
| *Aizoon canariense* | 0.008 | 0.024 | 0.271 | 1.000 | 0.702 |

**Table S3.** ANOSIM results for the plant communities inhabiting **platforms** and **ravines** habitats along the distribution area of *G. intermedia*. The average contribution of species to the differentiation between plateaus and platforms plant communities (**Average**), is presented along with the standard deviation of these contributions (**sd**); the relative abundance of species (**Abund**); the cumulative contribution of species (**c.sum**); and the p-values (***p***) of the ANOSIM permutations, in this case 999. Species with no abundance or contribution to the communities have been removed from the table.

| **Species** | **Average** | **sd** | **Abund** | **c.sum** | **p** |
| --- | --- | --- | --- | --- | --- |
| *Patellifolia patellaris* | 0.172 | 0.057 | 5.008 | 0.190 | 0.001 |
| *Argyranthemum foeniculaceum* | 0.137 | 0.067 | 0.343 | 0.341 | 0.001 |
| *Neochamaelea pulvurulenta* | 0.095 | 0.059 | 0.000 | 0.447 | 0.001 |
| *Plocama pendula* | 0.092 | 0.078 | 0.000 | 0.549 | 0.001 |
| *Euphorbia balsamífera* | 0.088 | 0.079 | 1.949 | 0.647 | 0.873 |
| *Opuntia dillenii* | 0.061 | 0.064 | 0.000 | 0.714 | 0.193 |
| *Cenchrus ciliaris* | 0.054 | 0.085 | 0.804 | 0.774 | 0.180 |
| *Mesembryanthemum nodiflorum* | 0.039 | 0.055 | 1.288 | 0.817 | 0.581 |
| *Kleinia nerifolia* | 0.035 | 0.059 | 0.403 | 0.856 | 0.604 |
| *Limonium pectinatum* | 0.030 | 0.038 | 0.000 | 0.889 | 0.931 |
| *Astydamia latifolia* | 0.020 | 0.042 | 0.676 | 0.912 | 0.635 |
| *Aeonium sp* | 0.020 | 0.041 | 0.000 | 0.934 | 0.002 |
| *Schizogyne sericea* | 0.019 | 0.038 | 0.000 | 0.954 | 0.884 |
| *Volutaria canariensis* | 0.015 | 0.030 | 0.000 | 0.970 | 0.002 |
| *Zygophyllum fontanesii* | 0.010 | 0.031 | 0.318 | 0.982 | 0.734 |
| *Fagonia cretica* | 0.009 | 0.028 | 0.326 | 0.992 | 0.829 |
| *Aizoon canariense* | 0.008 | 0.023 | 0.271 | 1.000 | 0.737 |

**Table S4.** ANOSIM results for the plant communities inhabiting **plateaus** and **ravines** habitats along the distribution area of *G. intermedia*. The average contribution of species to the differentiation between plateaus and platforms plant communities (**Average**), is presented along with the standard deviation of these contributions (**sd**); the relative abundance of species (**Abund**); the cumulative contribution of species (**c.sum**); and the p-values (***p***) of the ANOSIM permutations, in this case 999. Species with no abundance or contribution to the communities have been removed from the table.

| **Species** | **Average** | **sd** | **Abund** | **c.sum** | **p** |
| --- | --- | --- | --- | --- | --- |
| *Argyranthemum foeniculaceum* | 0.108 | 0.042 | 0.439 | 0.145 | 0.011 |
| *Neochamaelea pulvurulenta* | 0.080 | 0.048 | 0.000 | 0.252 | 0.012 |
| *Plocama pendula* | 0.077 | 0.065 | 0.000 | 0.356 | 0.019 |
| *Euphorbia balsamifera* | 0.074 | 0.071 | 5.042 | 0.456 | 0.953 |
| *Limonium pectinatum* | 0.074 | 0.055 | 2.871 | 0.555 | 0.088 |
| *Frankenia ericifolia* | 0.071 | 0.062 | 2.530 | 0.650 | 0.019 |
| *Opuntia dillenii* | 0.063 | 0.058 | 1.918 | 0.735 | 0.266 |
| *Schizogyne sericea* | 0.046 | 0.052 | 1.568 | 0.796 | 0.169 |
| *Kleinia nerifolia* | 0.040 | 0.060 | 1.003 | 0.850 | 0.496 |
| *Reseda scoparia* | 0.037 | 0.047 | 1.296 | 0.900 | 0.133 |
| *Cenchrus ciliaris* | 0.032 | 0.066 | 0.000 | 0.944 | 0.649 |
| *Aeonium sp* | 0.017 | 0.034 | 0.000 | 0.966 | 0.484 |
| *Fagonia cretica* | 0.014 | 0.029 | 0.480 | 0.985 | 0.777 |
| *Volutaria canariensis* | 0.011 | 0.023 | 0.000 | 1.000 | 0.430 |

**Figure S1.**
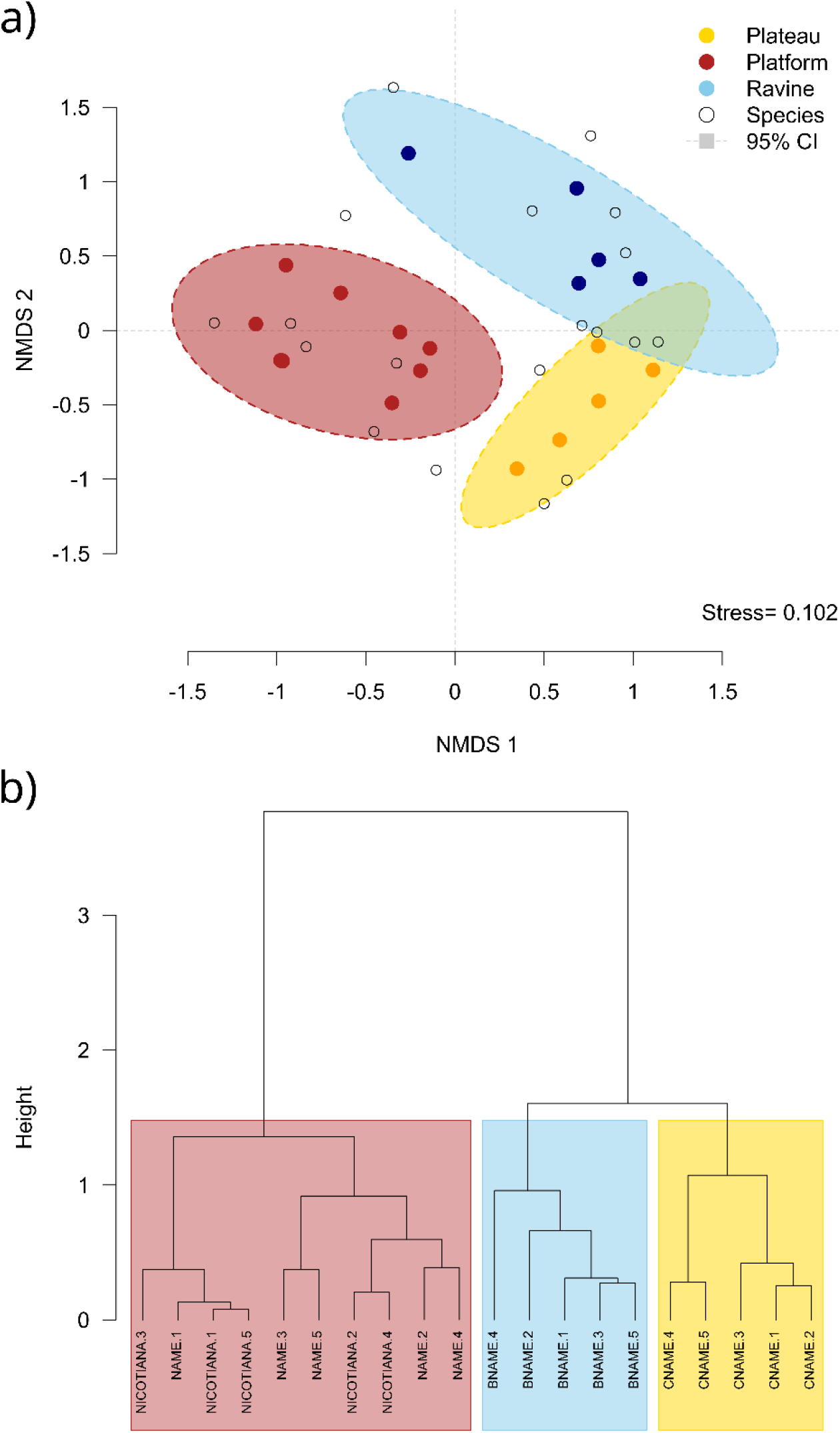
Non-metric multidimensional scaling (NMDS) of plant community composition for the different topographic features described within the distribution area of G. intermedia in the natural area of Guaza Mountain (**a**) and a hierarchical cluster of the vegetation communities (**b**). For the NMDS (a), each colour-filled point represents the random stratified sampling vegetation units, whereas the hollow circles represent the different species. Ellipses represent ordination 95% confidence intervals of the standard deviations. For the dendrogram (**b**), each leaf of the dendrogram represents a monitoring unit of the stratified random sampling of vegetation or a row of the Bray-Curtis dissimilarity matrix. Coloured boxes group sampling units according to their topographic features. Both in **a** and **b**, the different topographic features are marked with colours (red=Platform, blue=Ravine, and yellow=Plateau).

### S2. Delimitation of the breeding colony of yellow-legged seagulls (*Larus michahellis*) in the Natural Reserve of Guaza Mountain

To spatially extrapolate the results of the body-guard effect experiment to the rest of the distribution of *G. intermedia* that overlaps with *L. michahellis* breeding colony we first need to make a precise delimitation of such colony. Additionally, we need to predict the likely density of seagulls along its breeding area. For this, we combined on-site detection and delimitation along with remote sensing data and statistical modelling. To delimit the potential breeding and resting area of *L. michahellis*, we delimited 2 reference areas on which we confirmed the long-term presence of these animals. These reference areas were later projected onto a multi-channel high-definition aerial orthophotography (Figure S2.a-b). Using the range of pixel values from the Red, Green, and Blue (RGB) channels, we perform initial filtering of the image, selecting only the pixels that presented values on the three different channels for the areas where we have confirmed the presence of breeding seagulls (Figure S2.b). Based on this initial filtering we selected other two areas in which the satellite image confirms a strong concentration of seabird guano (Figure S2.b). Once we defined these areas of permanent presence of *L. michahellis*, we used a 2m resolution Digital Elevation Model (DEM) to extract relevant topographic features; Slope, calculated using the difference in height and orientation of the 4 adjacent pixels (Fleming & Hoffer, 1979; Ritter, 1987); Aspect, calculated using the (Fleming & Hoffer, 1979; Ritter, 1987); Ruggedness, calculated as the average angle of the 4 neighbouring pixels with respect to the horizontal plane (Fleming & Hoffer, 1979; Ritter, 1987); and distance to the sea, calculated as the minimum Euclidean distance between each pixel centroid and the sea (elevation equal to 0 in our DEM). Similar to the RBG channels previously described, we used the range of values from these terrain features to filter the spatial raster and select the areas more likely to be occupied by *L. michahellis* (Figure S2.C). To account for uncertainty and to remove gaps within the area classified as likely occupied by yellow-legged gulls, we added a 15 m buffer around it. Using high-definition orthophotography we confirmed that the selected areas matched the distribution of *L. michahellis* in the Natural Area of Guaza Mountain (Figure S2.e).

**Figure S2.**
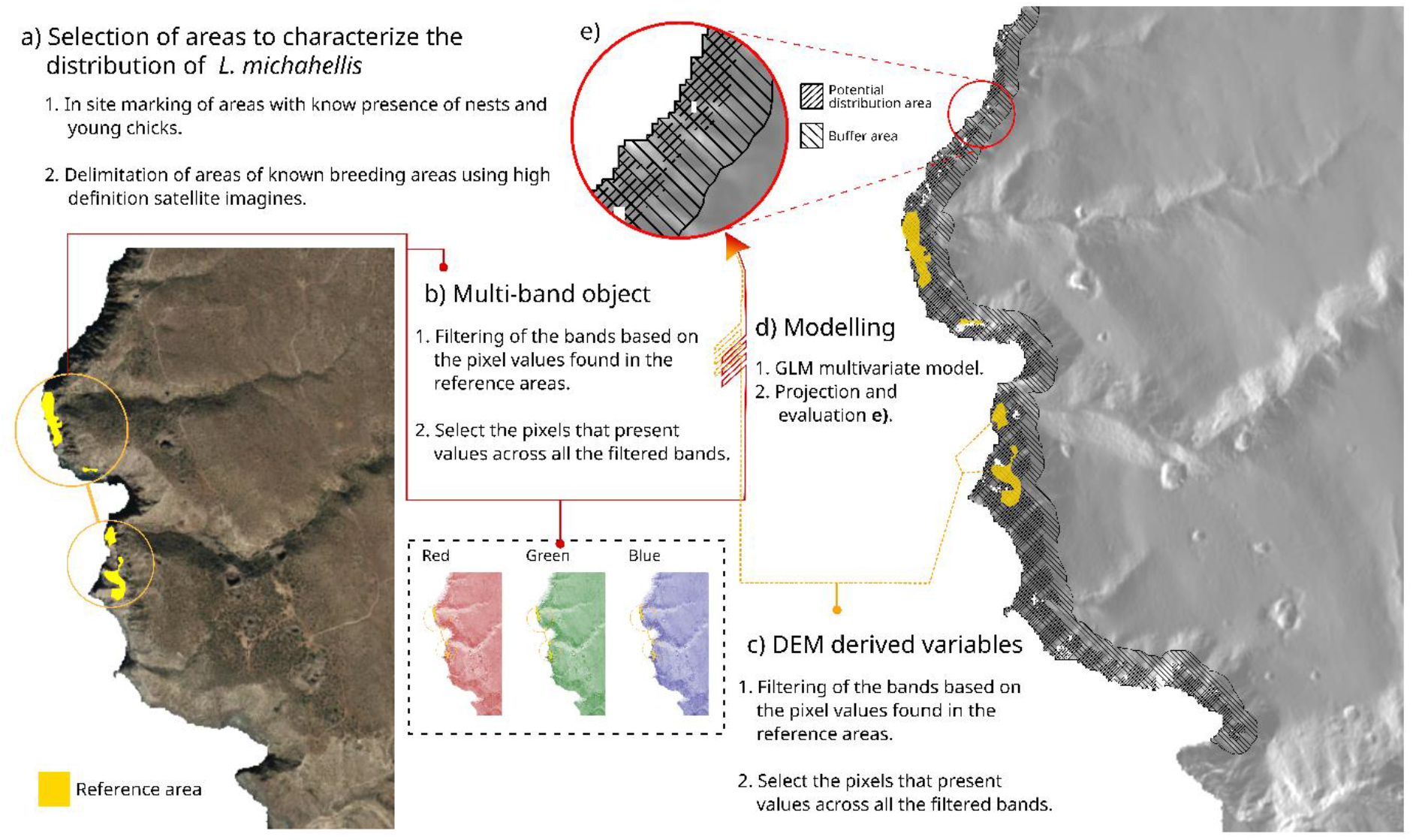
Workflow for the spatial delimitation of *Larus michahellis* breeding colony in the natural protected area of Guaza Mountain. The process implies the combination of field observations (counting of nest and seagull chicks around the sampling localities) and high-definition satellite data of areas occupied by *L. michahellis* (**a-b**). Once the areas of the known presence of the species are delimited, in situ, information about the gull’s density is mixed with precise digital topography variables to model the predicted density of *L. michahellis* chicks along the coastal cliffs (**c-e**).

To predict the number of young chicks across the potential distribution area of *L.michahellis*, first, we run a generalised linear model (GLM) with a Poisson distribution. For this model, we used the number of yellow-legged seagulls’ chicks around the points used for the body-guard effect experiment (see main text) as the response variable. For the explanatory variables we used the derived terrain features from the DEM, and averaged the values of all the pixels that fall into the 5 m radius of our counting points (Figure S2.c). To account for the differences in units between the different terrain variables we standardize the data by subtracting the mean and dividing the values by the standard deviation of the variables. The model showed that slope had a positive effect on the number of chicks whereas terrain aspect, distance to the sea, and terrain ruggedness had a negative effect. We use this fitted model to predict the potential number of chicks along the distribution area of *L. michahellis* breeding colony (Table S5). We use this new layer to make further predictions (see main text).

**Table S5.** Results from the GLM Poisson family were used to estimate the number of *Larus michahellis* chicks along the coastal cliffs of the natural protected area of Guaza Mountain. Term estimates, standard deviation and z-values from the models are presented along with the Wald-chi type II test performed to test the statistical significance of the terms on the response variable. Statistically significant terms are marked in bold.

| GLM family poisson model | | | | Wald-$\chi^2$ type II test | | |
| --- | --- | --- | --- | --- | --- | --- |
| Term | Estimated | sd | z-value | $\chi^2$ | d.f | p-value |
| intercept | -10.64 | 4.52 | -2.35 |  |  |  |
| Slope | 2.26 | 1.96 | 1.15 | 1.59 | 1 | 0.2 |
| Aspect | -0.62 | 1.41 | -0.43 | 0.18 | 1 | 0.66 |
| **Sea Dist** | **-7.7** | **3.57** | **-2.15** | **9.66** | **1** | **0.001** |
| Ruggedness | -2.37 | 1.79 | -1.32 | 2.12 | 1 | 0.14 |

Nagelkerke's R<sup>2</sup> 0.945

High-resolution orthophotography, as well as the high-definition DEM, were directly downloaded from the open repositories of the Spanish National Geographic Institute or ‘Instituto Geografico Nacional’ (IGN, https://www.ign.es/web/ign/portal) with the references ‘PNOA_MA_OF_REGC95_HU28_H50’ and ‘MDT02_REG95_HU28’ respectively. All the spatial analysis and terrain features calculations were performed in R using the Terra version 1.5-34 (Hijmans, 2022) and *raster* version 3.5-21 (Hijmans, 2022) packages. The GLM was fitted using the R-packages *stats* version 4.3.1 (R Core Team, 2021) and *car* version 3.1-2 (Fox & Weisberg, 2019).

### S3. Biometric data and exploratory analysis

**Table S6.**
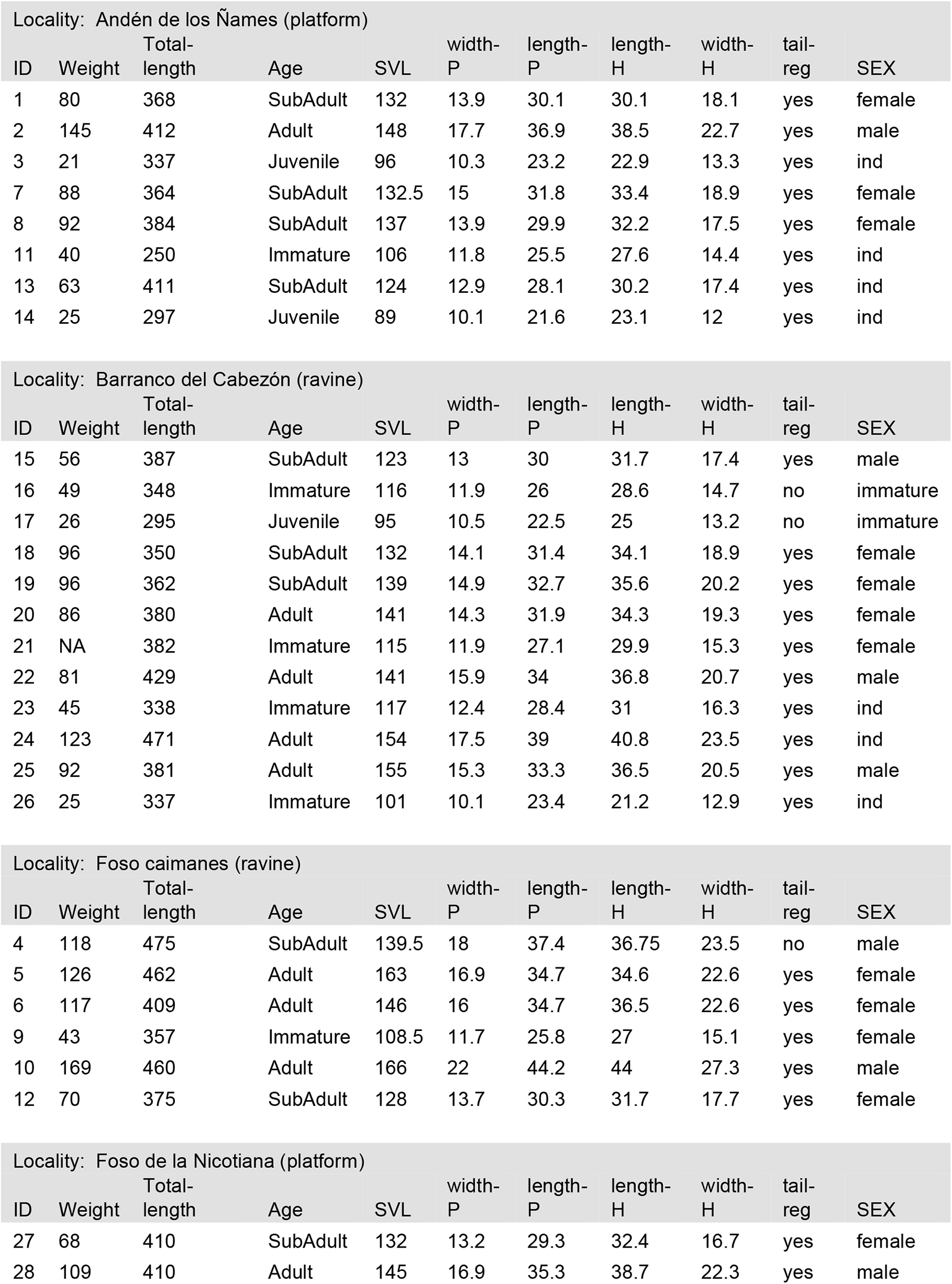
Biometric data of Gallotia intermedia by locality. The biometric data includes; Snout to Vent distance (SVL), measure ventrally in mm from the tip of the snout to the start of the cloaca; Pilleum length (Length-P), measure in mm following the total length of the pilleum scale complex; Pilleum width (widt-P), measure in mm along the widest section of the pilleum; Head length (length-H), mm from the back of the neck to the tip of the snout; Head width (width-H), mm across the widest portion of the head, normally around the cheeks; Total length, distance in mm from the tip of the snout to the tip of the tail; Age, classification according to the SVL of the animal (Adult= SVL>140; SubAdult= 120<SVL<140; Immature 100 < SVL <120; and Juvenile = SVL<100); Tail regeneration (tail-reg), binomial variable that take the value “yes” if the animals shows signs of recent or past tail regeneration, and “no” otherwise; Sex, sex of the animals (male, female, ind= adult individual that was not possible to determine their sex, immature= the animal was young at it was not possible to determine their sex by morphological traits).

| Locality: Andén de los Ñames (platform) |  |  |  |  |  |  |  |  |  |  |
| --- | --- | --- | --- | --- | --- | --- | --- | --- | --- | --- |
| ID | Weight | Total-length | Age | SVL | width-P | length-P | length-H | width-H | tail-reg | SEX |
| 1 | 80 | 368 | SubAdult | 132 | 13.9 | 30.1 | 30.1 | 18.1 | yes | female |
| 2 | 145 | 412 | Adult | 148 | 17.7 | 36.9 | 38.5 | 22.7 | yes | male |
| 3 | 21 | 337 | Juvenile | 96 | 10.3 | 23.2 | 22.9 | 13.3 | yes | ind |
| 7 | 88 | 364 | SubAdult | 132.5 | 15 | 31.8 | 33.4 | 18.9 | yes | female |
| 8 | 92 | 384 | SubAdult | 137 | 13.9 | 29.9 | 32.2 | 17.5 | yes | female |
| 11 | 40 | 250 | Immature | 106 | 11.8 | 25.5 | 27.6 | 14.4 | yes | ind |
| 13 | 63 | 411 | SubAdult | 124 | 12.9 | 28.1 | 30.2 | 17.4 | yes | ind |
| 14 | 25 | 297 | Juvenile | 89 | 10.1 | 21.6 | 23.1 | 12 | yes | ind |

| Locality: Barranco del Cabezón (ravine) |  |  |  |  |  |  |  |  |  |  |
| --- | --- | --- | --- | --- | --- | --- | --- | --- | --- | --- |
| ID | Weight | Total-length | Age | SVL | width-P | length-P | length-H | width-H | tail-reg | SEX |
| 15 | 56 | 387 | SubAdult | 123 | 13 | 30 | 31.7 | 17.4 | yes | male |
| 16 | 49 | 348 | Immature | 116 | 11.9 | 26 | 28.6 | 14.7 | no | immature |
| 17 | 26 | 295 | Juvenile | 95 | 10.5 | 22.5 | 25 | 13.2 | no | immature |
| 18 | 96 | 350 | SubAdult | 132 | 14.1 | 31.4 | 34.1 | 18.9 | yes | female |
| 19 | 96 | 362 | SubAdult | 139 | 14.9 | 32.7 | 35.6 | 20.2 | yes | female |
| 20 | 86 | 380 | Adult | 141 | 14.3 | 31.9 | 34.3 | 19.3 | yes | female |
| 21 | NA | 382 | Immature | 115 | 11.9 | 27.1 | 29.9 | 15.3 | yes | female |
| 22 | 81 | 429 | Adult | 141 | 15.9 | 34 | 36.8 | 20.7 | yes | male |
| 23 | 45 | 338 | Immature | 117 | 12.4 | 28.4 | 31 | 16.3 | yes | ind |
| 24 | 123 | 471 | Adult | 154 | 17.5 | 39 | 40.8 | 23.5 | yes | ind |
| 25 | 92 | 381 | Adult | 155 | 15.3 | 33.3 | 36.5 | 20.5 | yes | male |
| 26 | 25 | 337 | Immature | 101 | 10.1 | 23.4 | 21.2 | 12.9 | yes | ind |

| Locality: Foso caimanes (ravine) |  |  |  |  |  |  |  |  |  |  |
| --- | --- | --- | --- | --- | --- | --- | --- | --- | --- | --- |
| ID | Weight | Total-length | Age | SVL | width-P | length-P | length-H | width-H | tail-reg | SEX |
| 4 | 118 | 475 | SubAdult | 139.5 | 18 | 37.4 | 36.75 | 23.5 | no | male |
| 5 | 126 | 462 | Adult | 163 | 16.9 | 34.7 | 34.6 | 22.6 | yes | female |
| 6 | 117 | 409 | Adult | 146 | 16 | 34.7 | 36.5 | 22.6 | yes | female |
| 9 | 43 | 357 | Immature | 108.5 | 11.7 | 25.8 | 27 | 15.1 | yes | female |
| 10 | 169 | 460 | Adult | 166 | 22 | 44.2 | 44 | 27.3 | yes | male |
| 12 | 70 | 375 | SubAdult | 128 | 13.7 | 30.3 | 31.7 | 17.7 | yes | female |

| Locality: Foso de la Nicotiana (platform) |  |  |  |  |  |  |  |  |  |  |
| --- | --- | --- | --- | --- | --- | --- | --- | --- | --- | --- |
| ID | Weight | Total-length | Age | SVL | width-P | length-P | length-H | width-H | tail-reg | SEX |
| 27 | 68 | 410 | SubAdult | 132 | 13.2 | 29.3 | 32.4 | 16.7 | yes | female |
| 28 | 109 | 410 | Adult | 145 | 16.9 | 35.3 | 38.7 | 22.3 | yes | male |

### S4. Stable Isotopes Values

**Table S6.** δ^15^N and δ^13^C stable isotope values of vegetation collected at the Guaza Mountain at the cliff platforms, ravines and plateau habitats.

| Species | $\delta^{15}\text{N}$ | $\delta^{13}\text{C}$ | Foraging group | Family | Metabolic |
| --- | --- | --- | --- | --- | --- |
| *Argyranthemum spp.* | 4.14 | -27.68 | Ravine/Plateau | Asteraceae | C<sub>3</sub> |
| *Argyranthemum spp.* | 6.05 | -24.27 | Ravine/Plateau | Asteraceae | C<sub>3</sub> |
| *Argyranthemum spp.* | 7.14 | -22.65 | Ravine/Plateau | Asteraceae | C<sub>3</sub> |
| *Argyranthemum spp.* | 7.26 | -26.36 | Ravine/Plateau | Asteraceae | C<sub>3</sub> |
| *Argyranthemum spp.* | 7.90 | -24.15 | Ravine/Plateau | Asteraceae | C<sub>3</sub> |
| *Argyranthemum spp.* | 8.28 | -27.00 | Ravine/Plateau | Asteraceae | C<sub>3</sub> |
| *E. balsamifera* | 5.69 | -25.37 | Ravine/Plateu | Euphorbiaceae | C<sub>3</sub> |
| *E. balsamifera* | 5.90 | -25.73 | Ravine/Plateu | Euphorbiaceae | C<sub>3</sub> |
| *E. balsamifera* | 7.18 | -26.06 | Ravine/Plateu | Euphorbiaceae | C<sub>3</sub> |
| *E. balsamifera* | 7.41 | -25.82 | Ravine/Plateu | Euphorbiaceae | C<sub>3</sub> |
| *E. balsamifera* | 7.71 | -26.49 | Ravine/Plateu | Euphorbiaceae | C<sub>3</sub> |
| *E. balsamifera* | 9.32 | -24.58 | Ravine/Plateu | Euphorbiaceae | C<sub>3</sub> |
| *E. balsamifera* | 9.58 | -21.63 | Ravine/Plateu | Euphorbiaceae | C<sub>3</sub> |
| *E. balsamifera* | 9.88 | -21.74 | Ravine/Plateu | Euphorbiaceae | C<sub>3</sub> |
| *E. balsamifera* | 11.24 | -20.98 | Ravine/Plateu | Euphorbiaceae | C<sub>3</sub> |
| *E. balsamifera* | 11.38 | -20.95 | Ravine/Plateu | Euphorbiaceae | C<sub>3</sub> |
| *E. balsamifera* | 12.86 | -21.30 | Ravine/Plateu | Euphorbiaceae | C<sub>3</sub> |
| *E. balsamifera* | 12.96 | -19.89 | Platform | Euphorbiaceae | C<sub>3</sub> |
| *E. balsamifera* | 13.56 | -21.96 | Ravine/Plateu | Euphorbiaceae | C<sub>3</sub> |
| *E. balsamifera* | 13.95 | -21.30 | Ravine/Plateu | Euphorbiaceae | C<sub>3</sub> |
| *E. balsamifera* | 14.19 | -23.49 | Platform | Euphorbiaceae | C<sub>3</sub> |
| *E. balsamifera* | 14.37 | -21.04 | Ravine/Plateu | Euphorbiaceae | C<sub>3</sub> |
| *E. balsamifera* | 14.83 | -23.23 | Platform | Euphorbiaceae | C<sub>3</sub> |
| *E. balsamifera* | 14.91 | -21.23 | Platform | Euphorbiaceae | C<sub>3</sub> |
| *E. balsamifera* | 15.09 | -19.16 | Platform | Euphorbiaceae | C<sub>3</sub> |
| *E. balsamifera* | 15.40 | -21.05 | Ravine/Plateu | Euphorbiaceae | C<sub>3</sub> |
| *E. balsamifera* | 15.71 | -20.45 | Platform | Euphorbiaceae | C<sub>3</sub> |
| *E. balsamifera* | 17.71 | -23.54 | Platform | Euphorbiaceae | C<sub>3</sub> |
| *F. cretica* | 13.66 | -26.27 | Platform | Zygophyllaceae | C<sub>3</sub> |
| *F. cretica* | 15.21 | -24.95 | Platform | Zygophyllaceae | C<sub>3</sub> |
| *F. cretica* | 15.25 | -27.14 | Platform | Zygophyllaceae | C<sub>3</sub> |
| *F. cretica* | 15.39 | -27.41 | Platform | Zygophyllaceae | C<sub>3</sub> |
| *F. cretica* | 15.42 | -28.12 | Platform | Zygophyllaceae | C<sub>3</sub> |
| *F. ericifolia* | 7.65 | -22.33 | Ravine/Plateau | Frankeniaceae | C<sub>3</sub> |
| *F. ericifolia* | 7.79 | -23.02 | Ravine/Plateau | Frankeniaceae | C<sub>3</sub> |
| *F. ericifolia* | 8.33 | -24.02 | Ravine/Plateau | Frankeniaceae | C<sub>3</sub> |
| *K. nerrifolia* | 5.64 | -22.96 | Ravine/Plateau | Asteraceae | C<sub>3</sub> |
| *K. nerrifolia* | 6.11 | -20.94 | Ravine/Plateau | Asteraceae | C<sub>3</sub> |
| *K. nerrifolia* | 6.15 | -21.40 | Ravine/Plateau | Asteraceae | C<sub>3</sub> |
| *K. nerrifolia* | 12.63 | -20.62 | Platform | Asteraceae | C<sub>3</sub> |
| *K. nerrifolia* | 15.43 | -21.13 | Platform | Asteraceae | C<sub>3</sub> |
| *K. nerrifolia* | 15.59 | -20.24 | Platform | Asteraceae | C<sub>3</sub> |
| *L. intricatum* | 13.94 | -23.82 | Platform | Solanaceae | C<sub>3</sub> |
| *L. intricatum* | 15.07 | -25.70 | Platform | Solanaceae | C<sub>3</sub> |
| *L. intricatum* | 16.10 | -26.17 | Platform | Solanaceae | C<sub>3</sub> |
| *L. intricatum* | 16.54 | -25.09 | Platform | Solanaceae | C<sub>3</sub> |
| *L. intricatum* | 17.57 | -26.11 | Platform | Solanaceae | C<sub>3</sub> |
| *L. intricatum* | 18.26 | -27.18 | Platform | Solanaceae | C<sub>3</sub> |
| *L. intricatum* | 18.51 | -26.28 | Platform | Solanaceae | C<sub>3</sub> |
| *L. intricatum* | 19.15 | -27.75 | Platform | Solanaceae | C<sub>3</sub> |
| *M. crystallinum* | 14.49 | -16.74 | Platform | Aizoaceae | C<sub>4</sub>/CAM |
| *M. crystallinum* | 16.52 | -17.40 | Platform | Aizoaceae | C<sub>4</sub>/CAM |
| *M. crystallinum* | 17.61 | -15.47 | Platform | Aizoaceae | C<sub>4</sub>/CAM |
| *M. crystallinum* | 17.63 | -14.97 | Platform | Aizoaceae | C<sub>4</sub>/CAM |
| *M. crystallinum* | 18.16 | -14.59 | Platform | Aizoaceae | C<sub>4</sub>/CAM |
| *M. crystallinum* | 19.40 | -14.00 | Platform | Aizoaceae | C<sub>4</sub>/CAM |
| *N. glauca* | 15.44 | -23.93 | Platform | Solanaceae | C<sub>3</sub> |
| *O. dillenii* | 9.14 | -12.58 | Platform | Cactaceae | C<sub>4</sub>/CAM |
| *O. dillenii* | 11.41 | -12.81 | Platform | Cactaceae | C<sub>4</sub>/CAM |
| *O. dillenii* | 13.36 | -12.34 | Platform | Cactaceae | C<sub>4</sub>/CAM |
| *O. dillenii* | 14.87 | -13.99 | Platform | Cactaceae | C<sub>4</sub>/CAM |
| *P. patellaris* | 14.05 | -23.41 | Platform | Chenopodiaceae | C<sub>3</sub> |
| *P. patellaris* | 14.26 | -22.56 | Platform | Chenopodiaceae | C<sub>3</sub> |
| *P. patellaris* | 14.34 | -26.67 | Platform | Chenopodiaceae | C<sub>3</sub> |
| *P. patellaris* | 14.47 | -25.41 | Platform | Chenopodiaceae | C<sub>3</sub> |
| *P. patellaris* | 14.59 | -27.22 | Platform | Chenopodiaceae | C<sub>3</sub> |
| *P. patellaris* | 15.48 | -25.56 | Platform | Chenopodiaceae | C<sub>3</sub> |
| *P. patellaris* | 16.63 | -24.21 | Platform | Chenopodiaceae | C<sub>3</sub> |
| *P. patellaris* | 17.67 | -22.78 | Platform | Chenopodiaceae | C<sub>3</sub> |
| *P. patellaris* | 17.72 | -22.75 | Platform | Chenopodiaceae | C<sub>3</sub> |
| *P. patellaris* | 18.57 | -21.62 | Platform | Chenopodiaceae | C<sub>3</sub> |
| *P. patellaris* | 18.60 | -22.91 | Platform | Chenopodiaceae | C<sub>3</sub> |
| *P. patellaris* | 18.75 | -22.24 | Platform | Chenopodiaceae | C<sub>3</sub> |
| *P. pendula* | 4.38 | -25.39 | Ravine/Plateau | Rubiaceae | C<sub>3</sub> |
| *P. pendula* | 6.82 | -26.17 | Ravine/Plateau | Rubiaceae | C<sub>3</sub> |
| *P. pendula* | 8.11 | -27.07 | Ravine/Plateau | Rubiaceae | C<sub>3</sub> |
| *Portulaca spp.* | 16.69 | -14.79 | Platform | Portulacaceae | C<sub>4</sub>/CAM |
| *Portulaca spp.* | 19.49 | -14.00 | Platform | Portulacaceae | C<sub>4</sub>/CAM |
| *Portulaca spp.* | 20.12 | -14.24 | Platform | Portulacaceae | C<sub>4</sub>/CAM |

**Table S7:** *p-*values of pairwise comparisons of δ^13^C‰ and δ^15^N‰ values among forage item groups using a Wilcoxon rank-sum test. *: Statistically significant values p<0.05.

| **$\delta^{15}\text{N}$ values** | **Platform C<sub>3</sub>** | **Platform C<sub>4</sub>/CAM** | **Plateau/Ravine C<sub>3</sub>** |
| --- | --- | --- | --- |
| Group 1: Platform C<sub>3</sub> | - |  |  |
| Group 2: Platform C<sub>4</sub>/CAM | 0.51 | - |  |
| Group 3: Plateau/Ravine C<sub>3</sub> | 9.5e-14* | 3.7e-07* | - |
| Group 4: Seabirds-derived resources | 3.4e-10* | 0.00033* | 0.00847* |
| **$\delta^{13}\text{C}$ values** |  |  |  |
| Group 1: Platform C<sub>3</sub> | - |  |  |
| Group 2: Platform C<sub>4</sub>/CAM | 4.60e-11* | - |  |
| Group 3: Plateau/Ravine C<sub>3</sub> | 0.53 | 1.60e-10* | - |
| Group 4: Seabirds-derived resources | 4.70e-08* | 9.60e-07* | 1.30e-09* |

### S5. Media

**Figure S3.**
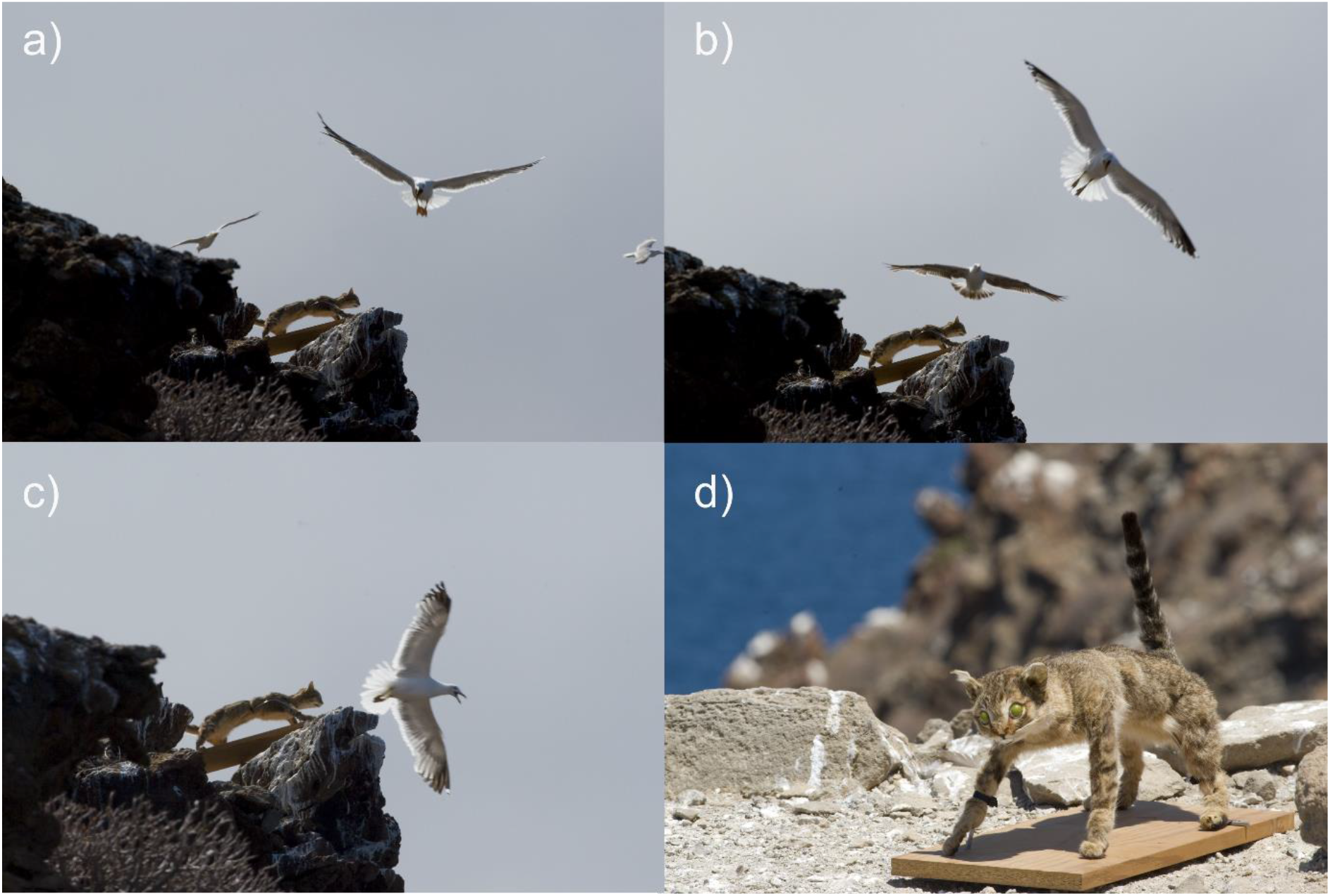
Exampled of recorded dive attack on the cat decoy (**a**-**c**) and cat decoy displayed in an open area near Guaza Mountain *L. michahellis* ground breeding colony (**d**).

**Figure S4.**
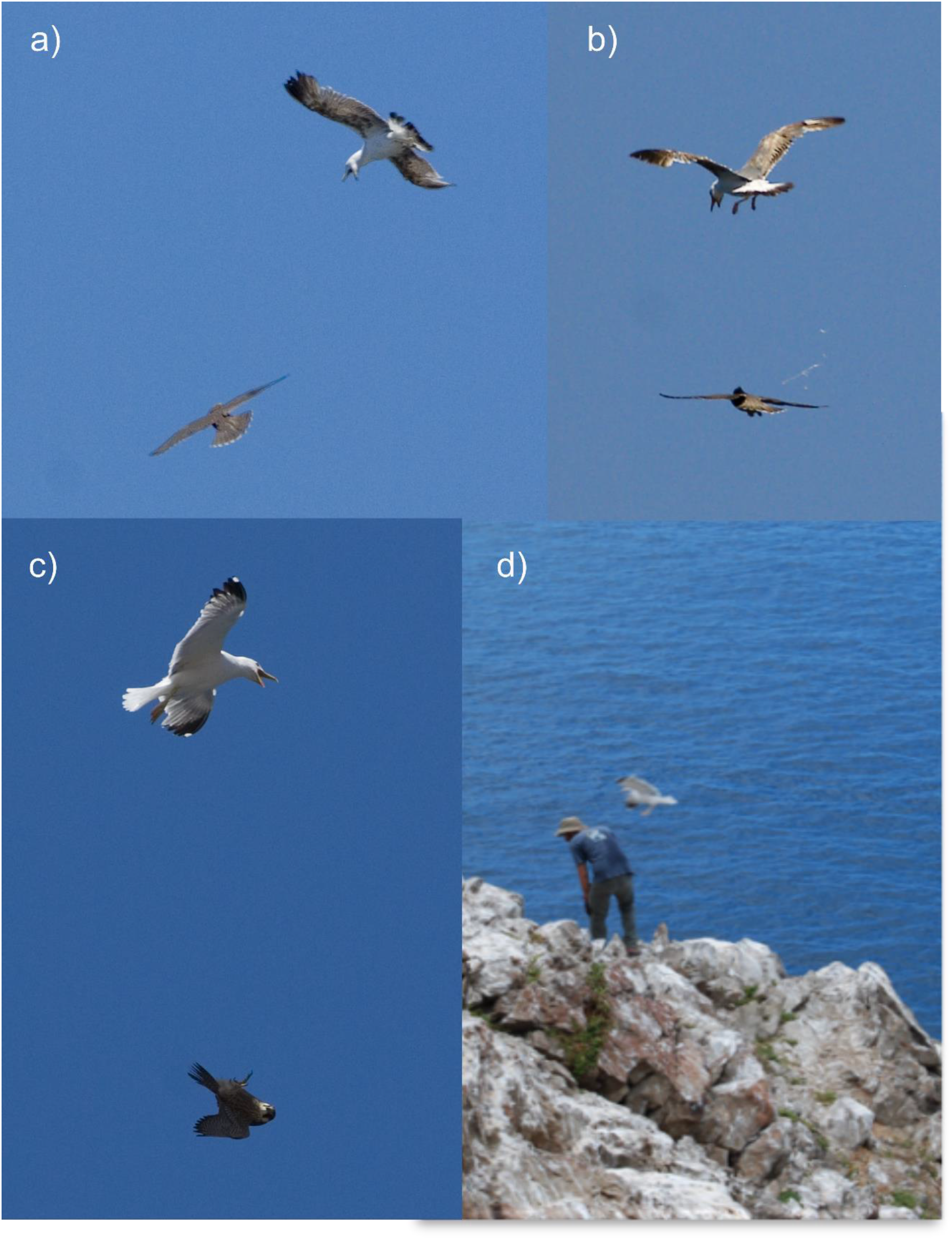
Exampled of recorded mobbing behaviour of *L. michahellis* over an Eleonora’s falcon over the breeding colony (**a**-**c**) and against one of the researchers while sampling into the breeding colony (**d**).

## Notes

### Competing Interest Statement

The authors have declared no competing interest.

### Summary of Updates

The changes included the correction of typos and general language problems found in the manuscript. In addition, some figure captions have been corrected to reflect the results better.

